# Human-aided dispersal and population bottlenecks facilitate parasitism escape in the most invasive mosquito species

**DOI:** 10.1101/2023.02.20.529246

**Authors:** Maxime Girard, Edwige Martin, Laurent Vallon, Van Tran Van, Camille Da Silva Carvalho, Justine Sack, Zélia Bontemps, Julie BaltenNeck, Florence Colin, Pénélope Duval, Simon Malassigné, Ian Hennessee, Lucrecia Vizcaino, Yamila Romer, Nsa Dada, Khan Ly Huynh Kim, Trang Huynh Thi Thuy, Christophe Bellet, Gregory Lambert, Fara Nantenaina Raharimalala, Natapong Jupatanakul, Clement Goubert, Matthieu Boulesteix, Patrick Mavingui, Emmanuel Desouhant, Patricia Luis, Rémy Cazabet, Anne-Emmanuelle Hay, Claire Valiente Moro, Guillaume Minard

## Abstract

During biological invasion process, species encounter new environments and partially escape some ecological constraints they faced in their native range, while they face new ones. The Asian tiger mosquito *Aedes albopictus* is one of the most iconic invasive species introduced in every inhabited continent due to international trade. It has also been shown to be infected by a prevalent yet disregarded microbial entomoparasite *Ascogregarina taiwanensis*. In this study, we aimed at deciphering the factors that shape the global dynamics of *As. taiwanensis* infection in natural *Ae. albopictus* populations. We showed that *Ae. albopictus* populations are highly colonized by several parasite genotypes but recently introduced ones are escaping it. We further performed experiments based on the invasion process to explain such pattern. To that end, we hypothesized that (i) mosquito passive dispersal (*i.e.* human-aided egg transportation) may affect the parasite infectiveness, (ii) founder effects (*i.e.* population establishment by a small number of mosquitoes) may influence the parasite dynamics and (iii) unparasitized mosquitoes are more prompt to found new populations through active flight dispersal. The two first hypotheses were supported as we showed that parasite infection decreases over time when dry eggs are stored and that experimental increase in mosquitoes’ density improves the parasite horizontal transmission to larvae. Surprisingly, parasitized mosquitoes tend to be more active than their unparasitized relatives. Finally, this study highlights the importance of global trade as a driver of biological invasion of the most invasive arthropod vector species.

**Significance:** Global trade expansion has facilitated the introduction of invasive species such as the Asian tiger mosquito *Aedes albopictus*. Eventually, invasive species might escape their natural enemies and this phenomenon exemplifies their invasion success. In this study, we combined field observations and laboratory experiments to decipher the ecological consequences of the invasion process on the interaction dynamics between *Ae. albopictus* and its most prevalent natural parasite *As. taiwanensis*. We observed a decrease in parasitism in recently introduced populations and provide experimental evidence to explain how human-aided mosquito transportation and mosquito population bottlenecks were a burden for the parasite.

## Introduction

Despite repeated calls on actions to prevent drastic consequences of global change, our society is now facing many ecological challenges that could profoundly affect its future (1). The cumulative effects of biological invasion and human activities are a threat to biodiversity due to change in global species distributions but also species loss and drift in community composition related to parasite transmission or competitive exclusion (2–7). Introduced species tend to raise health, agricultural, goods and services costs that weaken the global economy (8,9). Finally, some invasive species are considered as a threat to human health due to their pathogenicity or their ability to transmit pathogens (10). Currently, 37% of the 66 most-studied invasive species are considered as a threat while uncertainties remain on others (11). During biological invasion, invaders may be accompanied by parasites that may originate from their native range or an intermediate location (12). Co-introduction of invasive species with their parasites can lead to native species infection also referred as host shifting when the infection concerns different host species or spillover when the infection concerns different host populations (13). Invasive species may also acquire the parasite from native species. Finally, following introduction, invasive species may also escape the parasite colonizing them in their native or intermediate areas (14). Those events could contribute to exacerbate the performance and invasion success of recently introduced host species.

Both stochastic and selective pressures may lead to parasite release (15). Subsampling of individuals in the native or intermediate area during the invasion process may lead to uninfected newly formed populations. This is particularly true when the parasite prevalence is low within the source population (16,17). Furthermore, unsuitable conditions during the invasion path or colonization process may also lead to differential loss of the parasite due to *e.g.* unsuitable transportation conditions, higher host resistance at the invasion front and non-permissive environmental conditions (18). Other studies suggest that demographic changes may also alter the parasite transmission among individuals at the invasion front (19,20). Indeed, new populations are often founded by a small number of individuals that may be less prompt to exchange parasite due to lower horizontal transmissions and parasite virulence (21). In some cases, lower inter-individual transmission due to low population densities is counter balanced by a plastic change in host permissiveness toward parasites named the Density Dependent Prophylaxis (DDP) (22). In such cases, the immune system is relaxed in conditions of low host densities and reciprocally upregulated in conditions of high host densities. It is therefore expected that parasitism release is more likely to be exemplified if reduction in population densities occurs in absence of DDP. Finally, parasitism release may be the result of a parasite-driven counter selection on the host propensity to disperse. As an example, the parasite *Ophryocystis elektroscirrha* decreases flight performance of the Monarch butterfly *Danaus plexippus* reducing the chance of infected individuals to migrate (23).

The Asian tiger mosquito *Aedes albopictus* is one of the most invasive species and poses threats for human health (24) due to its ability to replicate and transmit at least 19 viruses such as Chikungunya, Dengue or Zika, as well as human and animal filarial nematodes (25,26). Originating from South and East Asia, its distribution has rapidly increased all over the world since 1980’s (27). Many studies pointed out the strong influence of human-aided transportation on intermediate and long distance dispersal of desiccated mosquito eggs and, in a lesser extent, adults (27–33). Actually, *Ae. albopictus* mosquitoes are often transported as desiccated eggs on attractive containers (*e.g.* used tires and lucky bamboo) (28,34–39). Eggs hatch whenever the containers get flooded after being introduced in a new area. Under optimal conditions, previous studies have shown that *Ae. albopictus* eggs can survive a month-long desiccation period (40). However, due to harsh transports conditions and population subsampling, the demography of recently introduced mosquito populations is often small (41,42). Such founder effects involve genetic drift, genetic bottlenecks and reductions in population size at the invasion front; however, these populations were shown to experience rapid growth in subsequent years (43,44,42,45,46). The final step of the invasion process is the local spread of mosquitoes and is both mediated either *via* flight or *via* passive transportations again (42,47).

*Ae. albopictus* populations are highly infected by an Apicomplexan gregarine parasite belonging to the *Ascogregarina* genus (48). Various species of *Ascogregarina* are colonizing mosquitoes with host specificity (49). *As. taiwanensis* is specifically associated with *Ae. albopictus* even though rare spillover events toward non-host mosquito species were reported (50–54). Its biological cycle is synchronized with the mosquito development (55). Oocysts released outside the eggshell are ingested by first instar larvae in water containers and replicate into developing individuals until adults release them. This parasite can be horizontally transmitted either when adults die, defecate or emerge within larval habitats. *As. taiwanensis* may have strong ecological consequences for its mosquito host although it is considered as a weak parasite (56). It was shown to have detrimental effects on mosquito traits such as development and survivorship at high density or when food is limited (57–59). A previous study conducted in Florida (USA) suggested that *Ae. albopictus* may escape *As. taiwanensis* at the invasion front (60). The authors demonstrated a drastically reduced parasite prevalence in recently introduced mosquito populations (<3 years). To reinforce those preliminary observations, we have conducted a field study in mainland France showing a similar trend of parasitism release at the invasion front. To estimate whether stochastic selection of uninfected individuals was likely to occur, we have also tested older populations in the native range and intermediate locations. We showed that the parasite is one of the most prevalent and dominant members of the mosquito microbiota in settled populations among contrasted locations across the world. Considering that stochastic selection of uninfected individuals was unlikely, we have thus conducted hypothesis driven experiments to test factors that may explain the observed release from the parasite pressure during biological invasion.

Our experiments have been designed to answer three questions based on the mosquito invasion process: (i) Does long distance dissemination through human-aided egg transportation limit the parasite maintenance? (ii) Does the decrease in host density following introduction affect the parasite dissemination probability in absence of DDP? and (iii) Does the parasite limit mosquito local active dissemination through flight? To answer those questions, we first tested the effect of desiccation on the parasite maintenance to reflect conditions occurring during long distance dissemination. Then, we tried to reproduce the decrease in mosquito populations reflecting founder effect through two different experiments to assess the impact of both adult and larvae densities on the parasite infection success. Finally, the impact of the parasitism on mosquito flight has been assessed through mosquito movement counting. Combined with previously published data, our results support that population density decrease following founding events and human-aided transportation contribute both to parasitism escape. Furthermore, our results suggest that the parasite *de novo* colonizes the mosquito with high densities two years after introduction.

## Material and methods

### Rearing parasitized and unparasitized mosquito lines

A total of ∼100 larvae were collected in 2017 in Villeurbanne (N: 45°46’18990’’ E: 4°53’24615’’) and Pierre-Bénite (N: 45°42’11534’’ E: 4°49’28743’’) in France. This population named AealbVB was grown into larger populations (∼5000 individuals) for 15 generations in a Biosafety level 2 insectary at 28°C, with a relative humidity of 80% and a day/night cycle of 18h/6h. Once the eggs were laid, they were immediately placed into water and ∼100 adults individuals were crushed and spread to make sure that the parasite was loaded. The water was never changed until the individuals molt into the 2^nd^ instar since individuals get infected during the 1^st^ instar. This protocol pursued for each generation. Uninfected individuals originate from the same initial population. To ensure that the parasite did not infect them, several actions were performed to shunt the infection path. First, the eggs were not immersed immediately but water was removed and eggs were kept dry at least 7 days before being flooded again to avoid direct transmission of this parasite when it is released by adults in water. Then, no contact was permitted between infected individuals and the water in which larvae were reared to avoid any direct transmission from feces or adult bodies. Finally, as the infection occurs during the first stage of larval development, the water was changed during the first instar to reduce the transmission probability. The absence of infection in the unparasitized population and presence of infection in the parasitized population was regularly checked by crushing adult individuals in water and observing the mixture at a magnification of x400 on an optical microscope (Leica).

### Sample collection and prevalence estimation

Live mosquito samples were collected with aspirators, net and BG-traps in 7 countries and 17 sites. They were identified using morphological characteristics (61), stored in 70% ethanol during transportation and frozen until use. Information related to site, country, year of collection, sex, climate, GPS coordinates and the population age (*i.e.* time between population introduction and sample collections) are reported in **Table S1**. This last information was obtained from local entomological surveillance services in France (EID and EIRAD), as well as from published reports in Italy (62), Spain (63) and USA (64). The date of introduction in Madagascar is relatively uncertain and was estimated to be older than 1904 (65). Populations from Thailand and Vietnam were considered as natives. DNA was extracted from surface sterilized mosquito whole bodies and PCR diagnostic were performed to determine the parasite prevalence (see **Supplementary material section**). To complete those dataset, supplemental data of *As. taiwanensis* presence-absence were extracted from previously published studies corresponding to mosquitoes collected in the USA (60,66,57,67) either from the manuscript or from graph by using the ImageJ software v.1.54f (68). Data are available on **Table S2**.

### Metabarcoding of eukaryotic and prokaryotic microbial communities

Using DNA extracted from surface sterilized mosquitoes, prokaryotic and eukaryotic communities within entire individuals were characterized through high throughput sequencing of hypervariable regions from the 16S rDNA and 18S rDNA genes respectively. The V5-V6 region of 16S rDNA gene was amplified using the primers 784F (5’-AGG ATT AGA TAC CCT GGT A-3’) and 1061R (5’-CRR CAC GAG CTG ACG AC-3’). The V1-V2 region of the 18S rDNA was amplified using the primers Euk82F (5′ GAA ACT GCG AAT GGC TC 3’) and Euk516R (5′ ACC AGA CTT GCC CTC C 3’). The details concerning PCR cycles are presented in supplementary materials. The PCR products were then sent to Biofidal sequencing company for purification and 2×300bp Miseq sequencing (Illumina). A total of 12,836,921 and 14,248,382 reads were obtained from 16S and 18S rDNA respectively and were demultiplexed using the Mothur pipeline (69) that is described in supplementary materials. After removing singletons and subsampling, 5,886 and 5,264 OTUs were retrieved respectively for 16S rDNA and 18S rDNA. Contaminant OTUs were removed if their proportion in samples were not at least 10-fold in comparison with controls. Finally, after quality control, data were homogenized by sampling 1,000 and 500 reads for 16S rDNA and 18S rDNA, respectively. The data have finally been converted into relative abundance, hereafter define as the proportion of reads associated to each OTU in comparison to the total number of reads per sample. Since relative abundances are of compositional nature, we have randomly selected 7 samples from each sex and group of microbial similarities for which we conducted 16S and 18S rDNA qPCR to convert relative abundances into absolute abundances *i.e.* number of copies associated with the given OTU per mosquito sample (see more details in **Supplementary materials**). Miseq sequences have been deposited on Zenodo under the project number 8252320 (https://zenodo.org/record/8252320).

### Testing the ability of parasite maintenance during passive long-distance transportation on desiccated eggs

As previously described, desiccated mosquito eggs can be transported by human transportations during weeks. However, we do not know if such transportation could affect the parasite. Since *Ae. albopictus* tends to prefer dark container (70), we let infected mosquitoes lay eggs on water containers coated with a dark green blotting paper and preserved them in the dark under optimal conditions (*i.e.* 28°C with a relative humidity of 80%). After respectively 35h, 1 week and 1 month, 100 eggs were harvested from the blotting paper and placed in a water container to hatch. Larvae were reared at 28°C and fed 1/3 yeast (Biovers) and 2/3 fish food and was provided *ad libitum* to the larvae. The adults that emerged from those containers were collected. The number of parasites colonizing the adult mosquitoes was estimated with qPCR on 10 males and females per time point and replicate. Primers and protocol for *As. taiwanensis* specific qPCR were developed for this study to determine the absolute abundance of the parasite (*i.e.* number of the parasite 18S copy per mosquito) and are detailed in the **supplementary material**. The experiment was reproduced five times. This experiment was repeated five times on five different blotting papers.

### Testing the influence of mosquito densities on inter-individual transmission

Water containers were maintained with 50ml of sterile water in cages containing either 100 (low density) or 3000 (high density) infected adult individuals during a week to allow them to release oocysts in water. The experiment was replicated 5 times for each condition (*i.e.* 5 donor mosquito cages were used for each tested mosquito density). Eggs from an unparasitized population were hatched in sterile water and 50 larvae were added to the container. Larvae were reared in similar conditions to the previously described experiment. Adult males and females were collected at emergence and their parasite density was measured with qPCR using 4 to 10 individuals per sex and replicate.

### Testing Density Dependent Prophylaxis (DDP)

To test whether the inter-individual transmission could be related to the density of larvae into the breeding sites, we hatched > 1000 larvae in containers to let them get infected with the parasite released from their parents. After 2 days, we then harvested them and grew them in new containers filled with distilled water at different densities (either 20, 50 or 100 larvae per 100ml of water). This protocol was replicated 5 times for each condition. Larvae were reared on a shelve providing shadow. When adults emerged, they were collected and parasites quantification was performed with qPCR (5 males and 5 females per replicate and larval density). For technical reasons, highest density modality could not be performed for the 5^th^ replicate (*i.e.* poor hatching rate for this replicate). Larval rearing conditions were similar to those in the previous experiment.

### Testing the influence of the parasite on active local mosquito dispersal through flight

Absence of the parasite in recently introduced mosquito populations might be the result of a poor propensity of infected mosquitoes to be active and fly. To test such hypothesis, parasitized and unparasitized adults were collected 2 weeks after their emergence and each individual was placed in a 25mm Ø activity tube containing a cotton soaked with a 10% sucrose solution and sealed with cotton. The tube was placed in a LAM25 Locomotor Activity Monitor system (Trikinetics). The system was crossed by 9 infrared beams that register an activity signal whenever the mosquito crosses the beams. Mosquitoes were left for 15min to rest before their activity was measured. The system was placed on a shelve that provides shadow to the mosquitoes since they tend to prefer such habitats in nature (71). The total activity measured with LAM systems can be mostly attributed to flight since walk distance do not exceed 20% of mosquitoes activity (72). Two experiments each involving 16 parasitized males, 16 parasitized females, 16 unparasitized males and 16 unparasitized females were conducted for 24h in a BSL2 insectary with 18h/6h day/night and 28°C. Whenever individuals died before the end of the registration, they were removed from the final dataset.

### Statistical and data analysis

#### Variations of As. taiwanensis prevalence in field populations

Statistical analyses performed with the R software v.4.3.1 (73). The proportion of parasitized mosquitoes (defined hereafter as parasite prevalence) has been modeled for field populations as a response variable using a binomial distribution and a probit link function. The explanatory variables tested were the sex and/or the mosquitoes collection site (defined hereafter as the mosquito origin), using the full dataset or a subsampled dataset for each sex to study the impact of factors without taking into account statistical interactions. The influence of each factor was tested with a Type II Wald χ^2^ test (*lme4* and *car* packages). We also correlated average parasite prevalence in collection sites with the number of years after the mosquito introduction as well as difference in prevalence between sites with the geographical distances separating them using the Spearman rank index with a correlation test and the Moran’s I autocorrelation coefficient respectively (*stat* and *ape* packages). Geographical distances between mosquito populations were calculated based on ellipsoid distance (*enmSdm,* package). Correlation analyses were repeated with data corresponding to previously published datasets from samples collected in the USA at different locations and time points after introduction.

#### Microbial communities associated with field populations

Microbiota diversity was estimated using the number of OTUs per individual sample after rarefaction for richness, Shannon index for alpha diversity and Bray-Curtis dissimilarity index for beta diversity (*vegan,* package). Richness and alpha diversities were modeled using a Poisson distribution and a log link function or a Gaussian distribution respectively and the impact of the sex and population origin were used as fixed factors. The fixed factors were tested with a type II Anova or Wald χ^2^ test (*lme4* and *car,* packages). The influence of sex and population origin on microbial Bray-Curtis dissimilarity were tested using permutational multivariate analysis (*vegan,* package). A similar analysis was used to compare the influence of the years after introduction and climates for each sex on the microbiota. Moreover, a mantel correlation test was used to compare the Beta diversity distances to geographical distances. Relative abundances of the 20 most dominant prokaryotic and eukaryotic OTUs were represented as a heatmap (*Complexheatmap,* package). The phylogenetic analysis of *As. taiwanensis* OTUs (based on an alignment of the most abundant sequences in each OTU) was performed with Seaview and the Muscle algorithm using other members of the Apicomplexa phylum. Phylogenetic trees were then generated with a Maximum Likelihood method, an HKY evolutionary model and 100 bootstraps (performed with Seaview and represented with Figtree). To identify the microorganisms which relative abundances covaried with the population age (years after introduction of *Ae. albopictus* on each sampling sites), a Constrained Analysis of Principal Coordinates (CAP) was performed (*vegan,* packages). The microbiota abundance correlation networks were performed with a homemade *python* code and visualized with the *pyvis* library. Briefly, the 200 most frequently retrieved OTUs for each community (prokaryotic and eukaryotic) were selected and their paired non-parametric correlations were tested. If the Spearman correlation (Rs) value was >0.25 (positive correlation) or <-0.25 (negative correlation), a linked was added between the two pairs. An automatic positioning algorithm was used based on mass-spring models. In this model, the links play a spring role which tend to bring the OTUs together while the OTUs play a role of opposite mass charges that are constantly repelled from each other. The algorithm used iterations to minimize the energy of the system. Individual *Ascogregarina taiwanensis* OTUs that vary according to mosquito population age were correlated with this last variable using non-linear Spearman correlations. Since relative abundances may induce bias due to their compositional nature, we have repeated CAP and correlation analyses with the selected samples for which we have performed qPCR.

#### Parasite prevalence, abundance and flight activity in controlled experiments

For the founder effect and desiccation experiments, the abundance of *As. taiwanensis* per mosquito were modeled with mixed models with a Gaussian and a Gamma inverse distribution respectively using mosquito sex, density or desiccation as fixed factors and the experimental replicates as random variables (*lme4*, package). Fixed factors were tested with an ANOVA or a type II Wald χ^2^ test. Post-hoc Tukey-HSD tests were used to estimate pairwise differences (*emmeans*, package). Prevalence was also tested for the experiment in which enough parasitized and unparasitized mosquitoes were detected (at least 5 events of each). It was analyzed with mixed model using a binomial distribution with a logistic link function with the mosquito sex, desiccation as fixed factors and tested with a type II Wald χ^2^ test. Flight activities (*i.e.*number of movements for each mosquitoes) was modeled with mixed model using a negative binomial, the sex of mosquitoes and infection status toward *As. taiwanensis* as fixed factors and the experimental replicates as random variables. The impact of the fixed factors was tested with a type II Wald χ^2^ test. The datasets and scripts used for statistical analysis have been deposited on zenodo (https://zenodo.org/record/8252320).

## Results

### Recently introduced mosquitoes in mainland France harbor lower prevalence of *As. taiwanensis*

The prevalence of *As. taiwanensis* was estimated in 598 mosquitoes sampled among 17 populations using diagnostic PCR (**Table S1**; **Fig. S1**). Samples collected in mainland France were a combination of recently introduced populations and intermediate ones. For all other countries, native or intermediate populations were collected. Among the studied variables, the population origin significantly impacted the prevalence of the parasite and differences were observed between sexes along the sampling sites (**Table 1**). For those reasons, the two sexes were then analyzed separately. To explain features of the mosquito origin that can influence the prevalence of the parasite, we further tested the impact of climate, population age and geographical distance between mosquito populations. Variations in climatic regions has no significant impact on parasite prevalence (**Table 1, Figure S2**). However, it was positively correlated with the mosquito population age for both males and females (**Table 1**, **Figure 1A, 1B**). In recent mosquito populations (< 2 years), the observed prevalence of *As. taiwanensis* ranged from 0 to 25%, while it ranged from 37% to 100% in populations that were native or introduced for more than 6 years. This effect was specifically strong on French sites that are geographically closed from each other but harbor high differences in parasite abundance correlating with differences in population ages. We could therefore assume that spatial autocorrelation might influence this observed pattern. The Moran’s index revealed a very weak but significant spatial autocorrelation for Mosquitoes populations at the global scale for both female and male mosquitoes (I = 0.27, p=0.038; I = 0.28, p = 0.013 for both sexes respectively). When considering the French populations separately no significant autocorrelation was evidenced for females (I = 0.05, p=0.064) and a very weak and yet significant autocorrelation was evidenced for males (I = 0.16, p = 0.02). To separate the impact of population age from that of geographical distances, we included prevalence data from previously published studies concerning recently introduced and older populations from the USA (**Table S2**). Those data referred to prevalence of *As. taiwanensis* in mosquito larvae and mixed adults and were included separately but in comparison with our study (**Figure 1B**). The spatial autocorrelation was not significant in this new dataset (I = −0.52, *p* = 0.22) while correlation between parasite prevalence and population age was comparable with the one reported for French populations (R_s_=0.46, *p*<0.001).

**Fig. 1.**
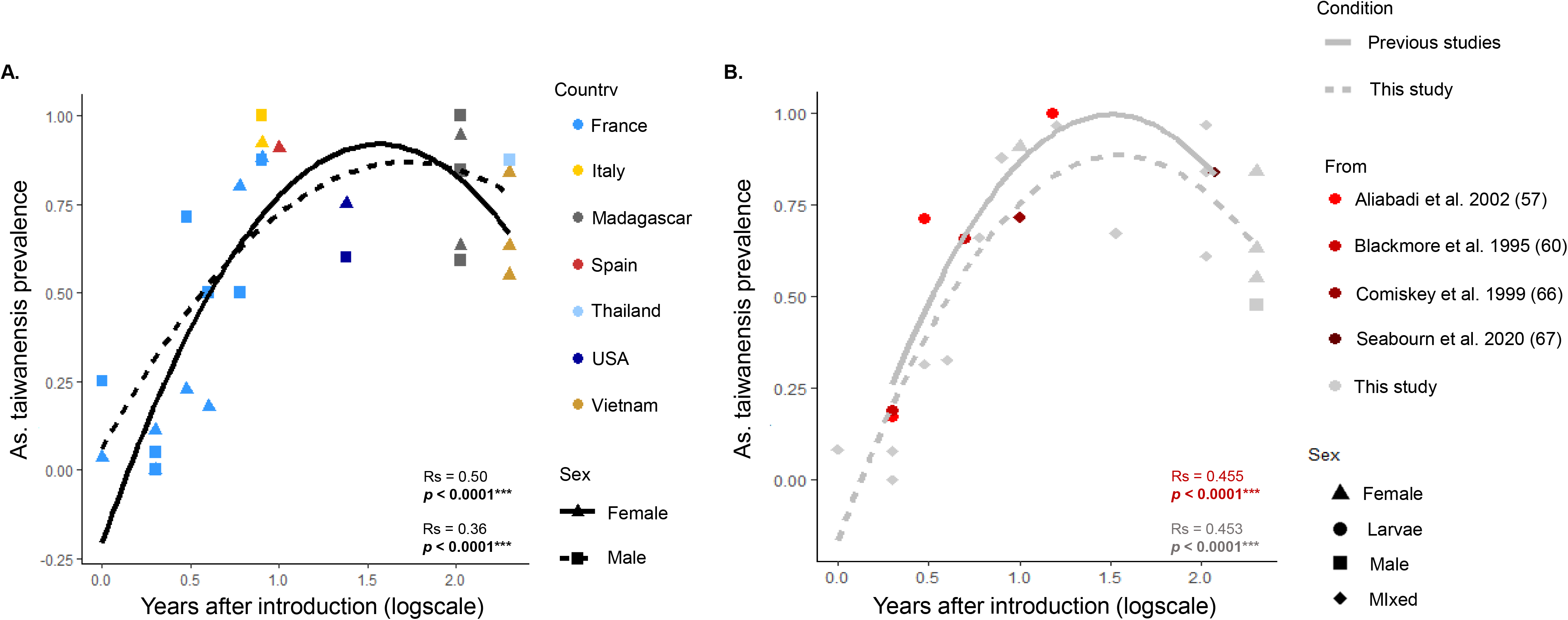
Correlation between the prevalence of *As. taiwanensis* and the time after introduction of the mosquito populations in each sampling site. (A) Correlation between parasite prevalence (*i.e.* proportion of infected individuals) and the population ages (number of years separating the mosquito introduction from the collection date) for females (triangle) and males (square) have been investigated in different countries (colors). (B) A comparative analysis has been realized between previously published data and the data from this study regarding the correlation between parasite prevalence and population ages. The data from previously published studies includes larvae (circles) and mixed adults (diamond) and are colored in red meanwhile the data from this study are presented in grey. Each dot represents a sampling site. Native populations were arbitrarily set at 200 years to be ranked at the end of the x-axis.

**Table 1.**
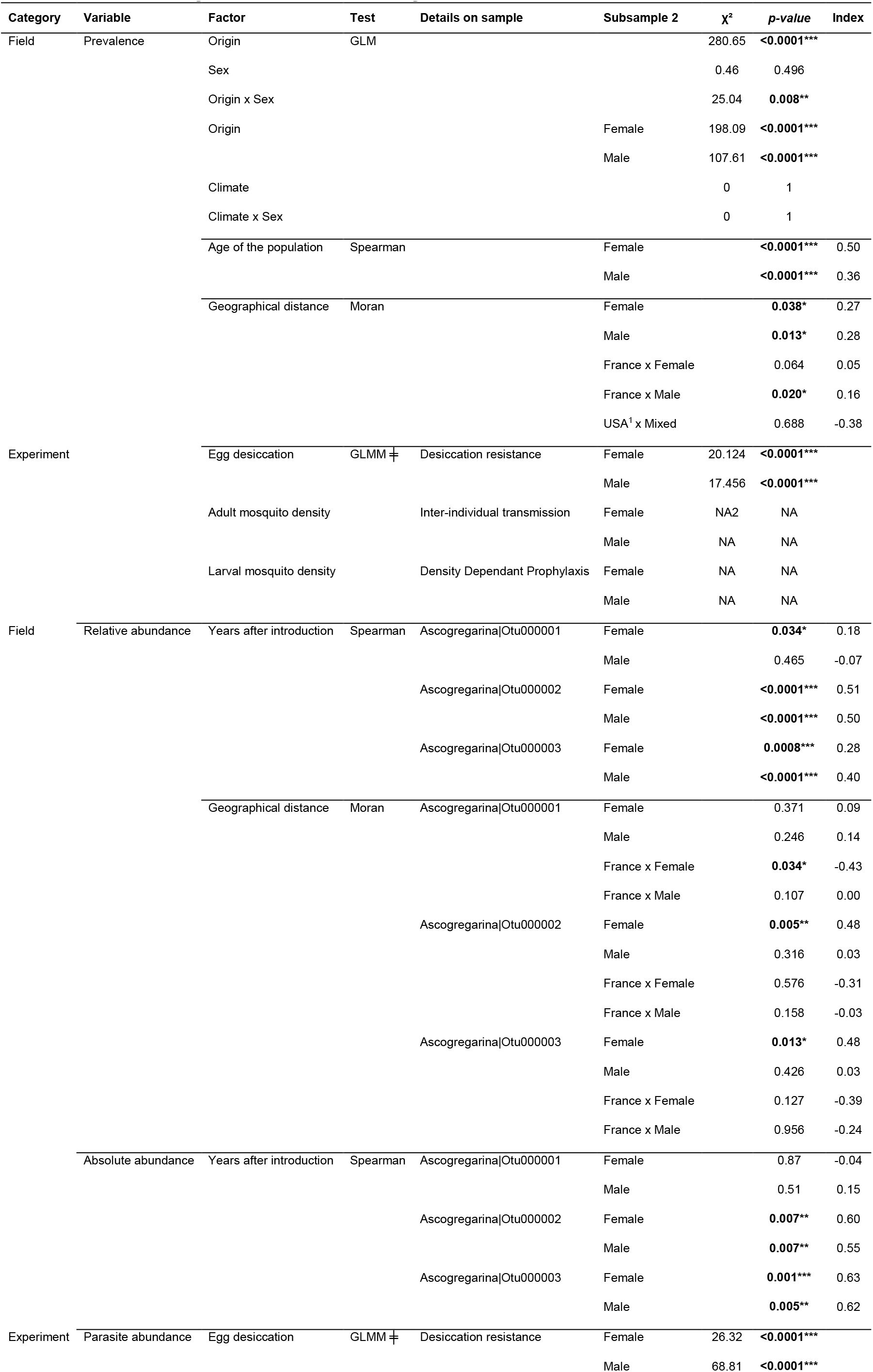
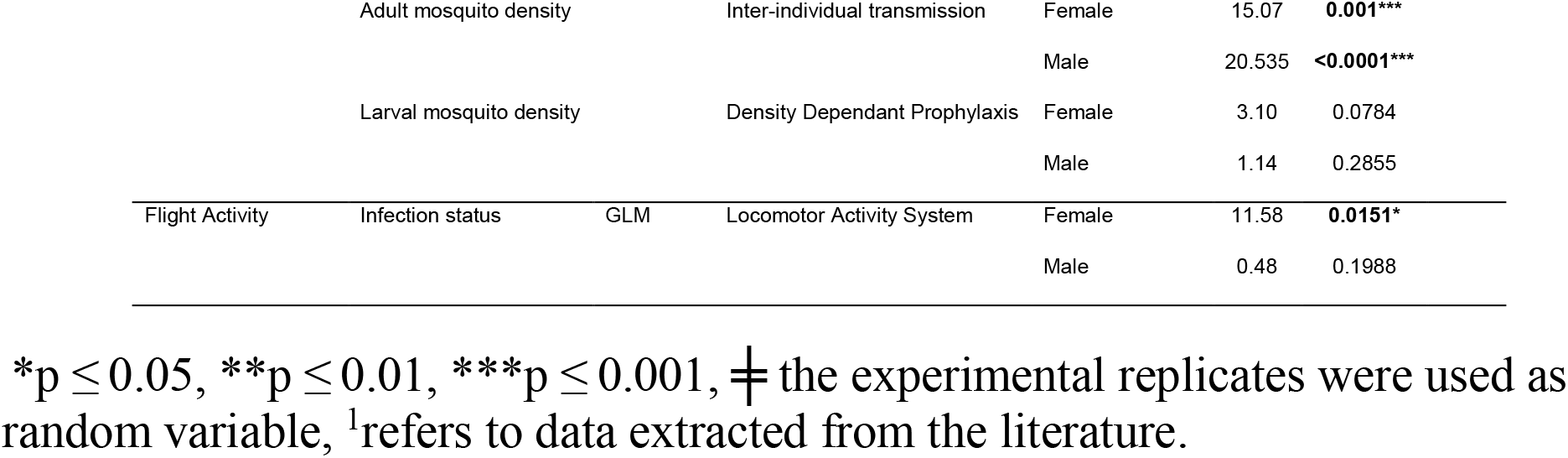
Summary of statistical analyses.

### Ascogregarina taiwanensis abundances correlate with the microbiota composition, the mosquito origin and the population age

To compare dynamics of *As. taiwanensis* within the microbiota of the different mosquito populations, a metabarcoding analysis was conducted on a subset of 248 male and female mosquitoes (**Table S1**). Mosquitoes were randomly selected for this analysis within the previously described populations. The eukaryotic and prokaryotic microbiota of *Ae. albopictus* were respectively dominated by two OTUs classified as *Wolbachia* (**Figure 2A**) and three OTUs classified as *Ascogregarina* (**Figure 2A**). In total 8 different OTUs of *Ascogregarina* were identified among the 200 most frequently retrieved OTUs but 5 showed low abundances. Those three dominant *Ascogregarina* OTUs showed a prevalence of 74.4% and an average abundance of 42.1±41.2% (mean±s.d.) among the eukaryotic microbiota while the two dominant *Wolbachia* OTUs showed a prevalence of 93.4% and an average abundance of 39.4±34.4% among the prokaryotic microbiota. Other OTUs identified as *Aureobasidium* and *Penicillium* genera for eukaryotes or as *Cutibacterium*, *Staphylococcus* and *Acinetobacter* genera for prokaryotes were also prevalent. Study of factors involved in variation of microbiota composition (*i.e.* β-diversity) between individuals revealed that the origin of mosquitoes was the most important (R^2^=0.32, *p*=0.001) structuring factor in interaction with the sex (R^2^=0.04, *p*=0.001) at a lower magnitude (see **Table S3**). Both eukaryotic and prokaryotic microbiota dissimilarities were also correlated with the age of the population (**Figure S2**), climate (**Figure S3**) and geographical distance between individuals (**Figure S4**).

**Fig. 2.**
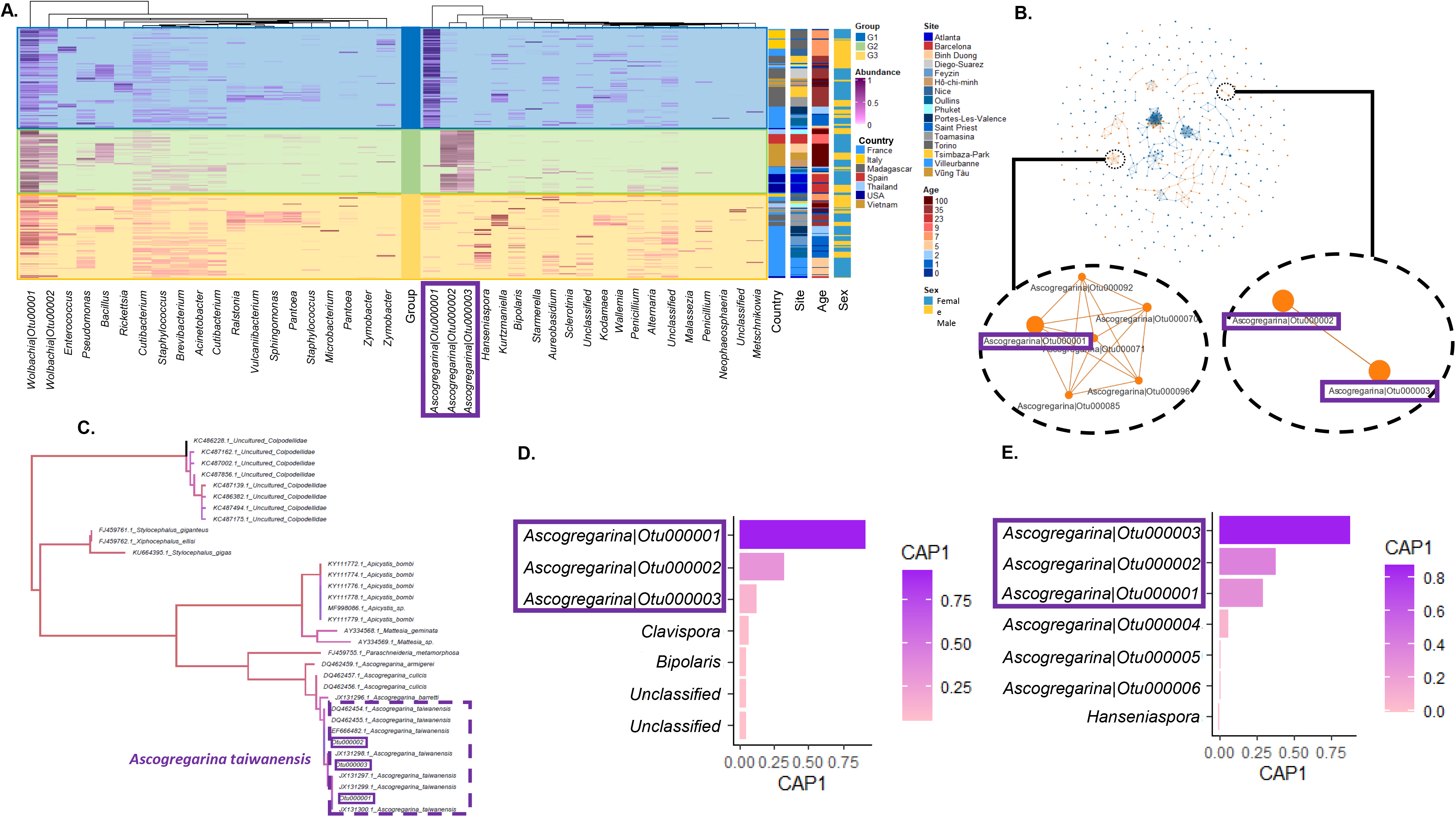
Heatmap of the microbiota associated with *Ae. albopictus*. (A) The 20 most abundant prokaryotic and eukaryotic OTUs were represented in columns. The number attached to *Wolbachia* and *Ascogregarina* OTUs corresponds to the OTUs number in the following analysis. The ‘Abundance’ color scale shows the OTU proportion in the community (pink). Rows are classified depending on the central scale that represent three different groups based on eukaryotic community dissimilarity (Bray-Curtis distance) and that correlates with variations in the three *As. taiwanensis* OTUs proportions. Individuals belonging to the G1 group are dominated by *Ascogregarina* Otu000001, those belonging to the G2 group are dominated by *Ascogregarina* Otu000002 and *Ascogregarina* Otu000003 and those belonging to the G3 group are not dominated by *Ascograrina* OTUs. Three additional scales show colors corresponding to the individuals’ origin site, age (years after introduction), and sex. (B) A network analysis showing positive correlations between microbial OTUs (eukaryotes are represented in yellow while prokaryotes are in blue and each link represents a correlation with ρ>0.25). The left network in the zoom is dominated by the *As taiwanensis* Otu000001 and is correlated with minor OTUs assigned to *Ascogregarina*. The right network in the zoom shows correlations between the *As. taiwanensis* Otu000002 and Otu000003. (C) Phylogenetic analysis revealed the belonging of OTU identified as Gregarinasina to the species *Ascogregarina taiwanensis*. (D) The main eukaryotic OTUs according to the coordinate of the axis (CAP1) extracted from a constrained analysis of principal coordinates (Capscale) on the OTU relative abundance highlighting correlations between OTUs and the age of the population. The three *As. taiwanensis* main OTUs are circled in violet. (E) The main eukaryotic OTUs according to the coordinate of the axis (CAP1) extracted from a constrained analysis of principal coordinates (Capscale) on the OTU absolute abundance highlighting correlations between OTUs and the age of the population. The three *As. taiwanensis* main OTUs are circled in violet.

Based on their similarity, the eukaryotic community could be separated in three groups (**Figure 2A**). The first group (60 individuals) was dominated by the most prevalent OTUs of *Ascogregarina* (Otu000001), the second (43 individuals) by two different OTUs of *Ascogregarina* (Otu000002 and Otu000003), and the last one (84 individuals) by neither of those three OTUs. This pattern was confirmed with correlations networks that showed two independent networks of positive correlations between relative abundances of OTUs identified as *Ascogregarina* (**Figure 2B**). The first network involved the dominant Ascogregarina (Otu000001) and fifth other underrepresented *Ascogregarina* OTUs (Otu000070, Otu000071, Otu000085, Otu000092, Otu000096) while the second one involved solely the two others dominant *Ascogregarina* OTUs (Otu000002 and Otu000003). The three dominant *Ascogregarina* OTUs could be confidently classified within the *As. taiwanensis* species based on a maximum likelihood phylogeny conducted on this same 18S rDNA region (**Figure 2C**). The three most abundant *Ascogregarina taiwanensis* OTUs were explaining most part of the eukaryotic microbiota that covaries with the age of the population based on a CAP analysis (**Figure 2D**). Correlations of relative abundances of those three OTUs showed that Otu000002 and Otu000003 increased in abundance in old populations while Otu000001 significant vary with the age of the population only for females with a low correlation index (**Figure 3A**). For each of the three group of microbiota similarity, absolute abundances were calculated for a subset of 7 females and 7 males. When converted in absolute abundances, the three most abundant *Ascogregarina taiwanensis* OTUs were still the most discriminant ones covarying with the age of the population in the CAP axis (**Figure 2E**). Otu000002 and Otu000003 showed a significant positive correlation between their absolute abundance and the age of the mosquitoes’ population while this was not significant for Otu000001 (**Figure 3B**).

**Fig. 3.**
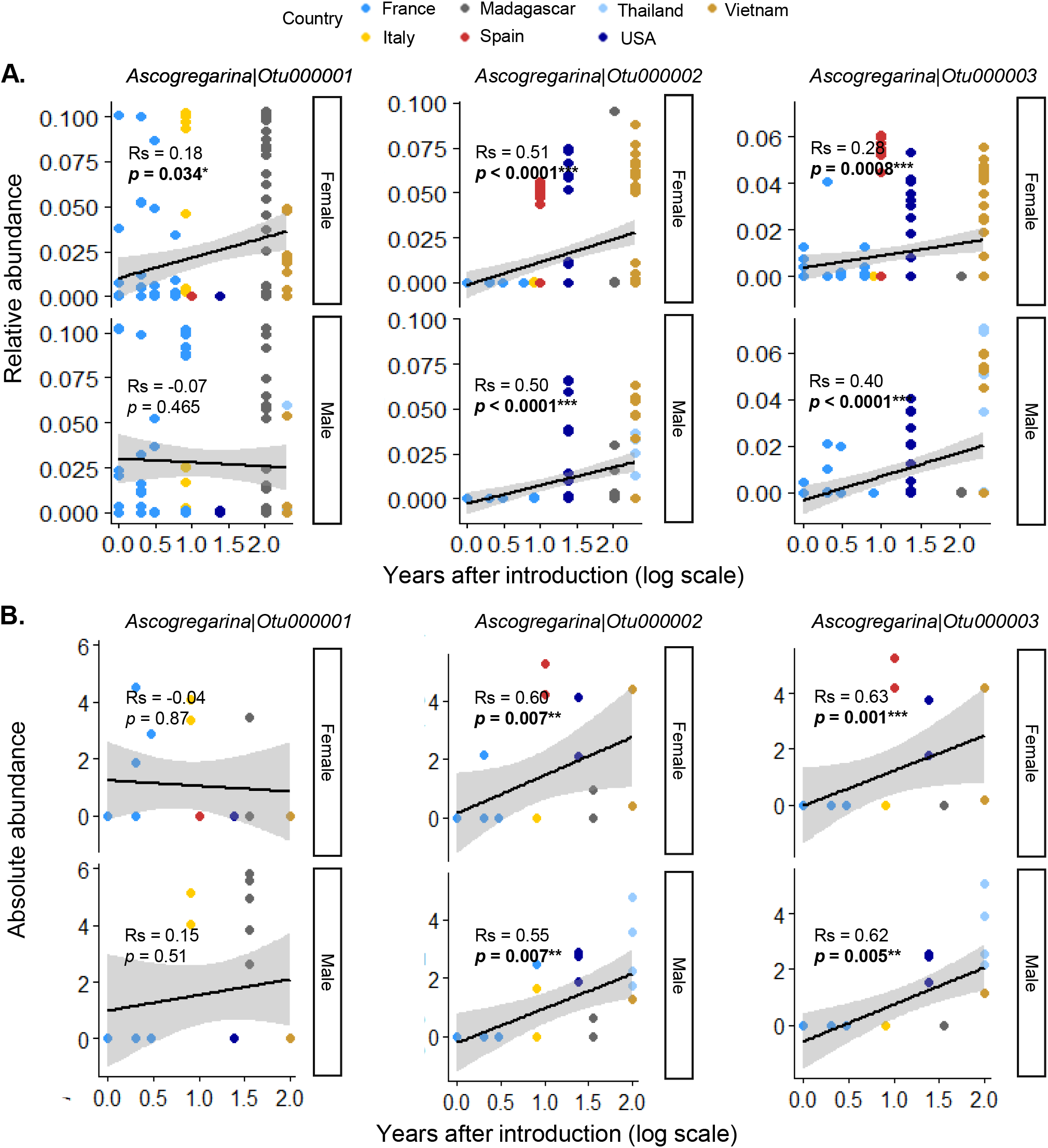
Correlations between *Ascogregarina* OTUs abundance and the mosquito population age. (A) The age of the mosquito populations and the relative abundance of the three dominant *Ascogregarina* are represented for female and male mosquitoes. (B) The age of the mosquito populations and the absolute abundance of the three dominant *Ascogregarina* are represented for female and male mosquitoes. The p-value represent the significance of the spearman correlation between OTUs absolute abundances and the age of the population (number of years between introduction and sampling). The native population was set to 200 years after introduction to be ranked at the end of the x-axis. The points are colored based on the mosquito population country.

### Parasitic success decreases over time when desiccated eggs are stored during human-aided transportation

To evaluate whether egg desiccation may restrain the parasite infectivity through long distance transportation, we collected eggs from five containers. Desiccated eggs were separated into three groups (100 eggs/group) that we respectively stored 35h, 1 week and 1 month before being flooded (**Figure 4A**). Parasite quantifications in adults were performed on 10 individuals per time point and replicate. Results showed significant decreases of the parasite absolute abundance, hereafter defined as the number of copies for specific genes of *As. taiwanensis*, over time after desiccation for both males and females (**Figure 4B**, **Table 1**). Based on 95% confidence interval of the model predictions, between 4421.4 and 86.6 copy of oocysts per ng of DNA were quantified per mosquito at the beginning of the experiment while this number was comprised between 134.3 and 0.2 after a month period of desiccation. Such treatment also affected the prevalence of the parasite within mosquitoes (**Figure 4C**, **Table 1**).

**Fig. 4.**
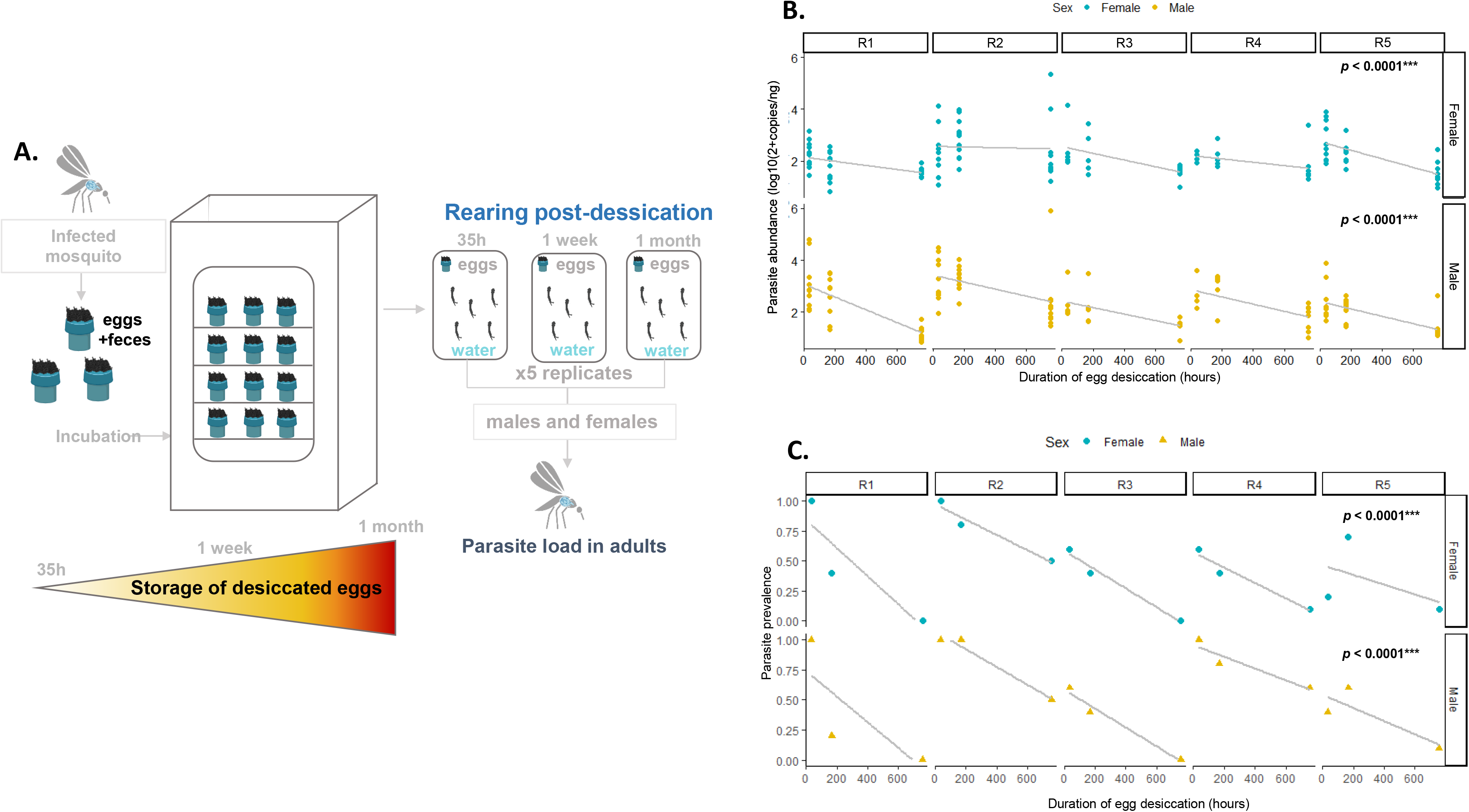
Impact of egg maintenance outside of water (desiccated) on the infectivity of *As. taiwanensis*. (A) Desiccated eggs have been stored for 35h, 1 week or 1 month in a container before being flooded to trigger egg hatching. Five containers were tested for each time point. The parasite abundance was estimated on emerging adults by qPCR. (B) The impact of desiccation time on parasite abundance was represented for each repeat (R1 to R5) and both sexes. (C) The impact of desiccation has also been tested on the parasite prevalence and was represented for each repeat (R1 to R5) and both sexes. The p-value and associated statistics have been obtained through spearman correlation analysis.

### Founder effects can reduce the amplitude of parasite infections due to low mosquito population densities and an absence of Density-Dependent Prophylaxis (DDP)

Two experiments were designed to determine whether the density of adults releasing oocysts and DDP (*i.e.* number of larvae receiving oocysts) could influence the parasite success to colonize new adult mosquitoes (**Figure 5A**). When the number of adults releasing oocysts in water was increased, we observed that higher densities of parasites (emmeans estimate ± se: 207±45.7 and 191±49.8 for males and females respectively) colonized individuals emerging from the water habitat (**Figure 5B**, **Table 1**). We also noticed that emerging females harbored more parasite oocysts than males (emmeans estimate ± se: 100±33.5). However, the parasite success was not significantly modified by larval densities either in males or in females (**Figure 5C**, **Table 1**).

**Fig. 5.**
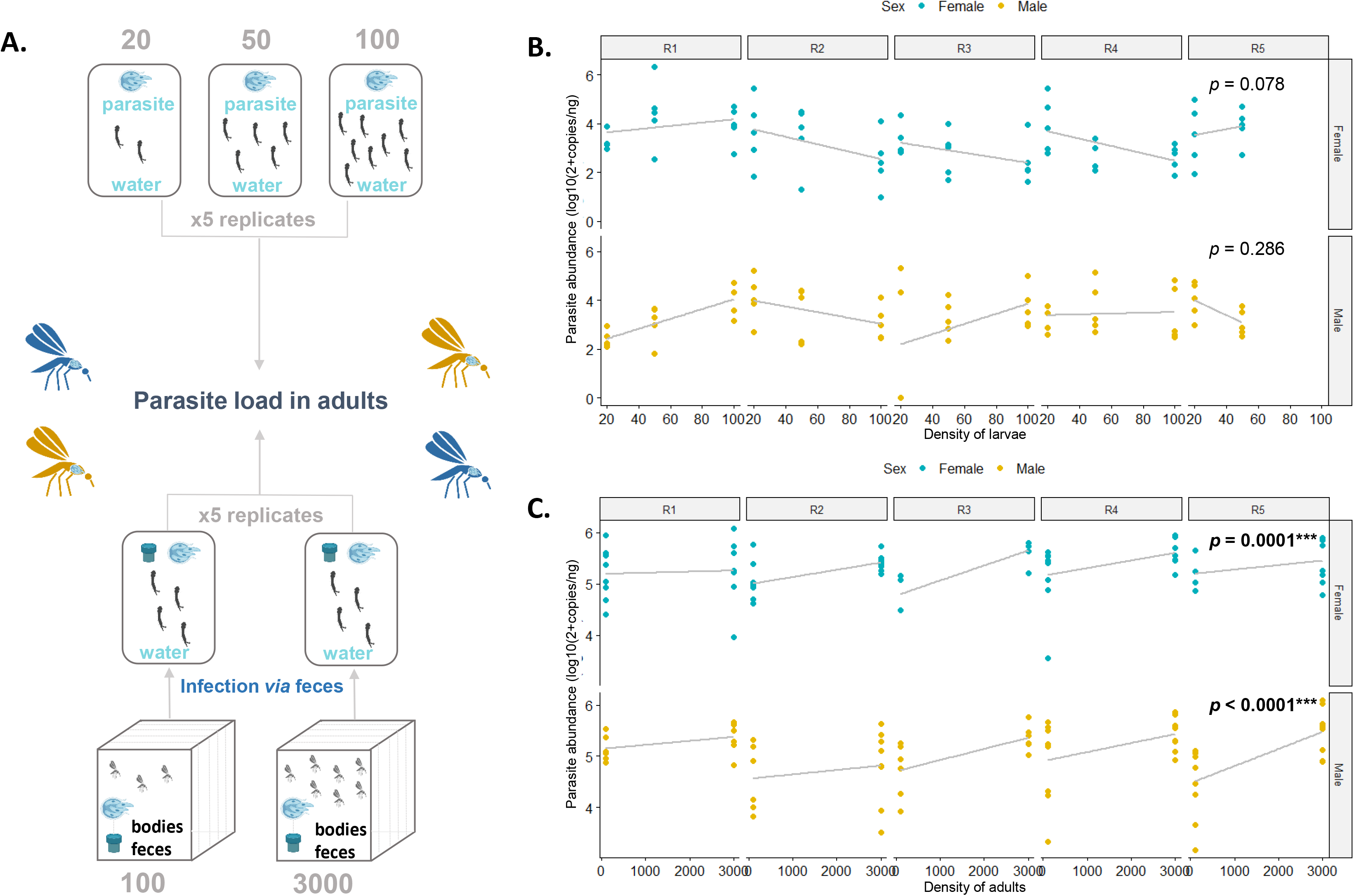
Density of adults releasing oocysts positively impacts the gregarine colonization and it is not counter balanced by density dependent prophylaxis. (A)The impact of adult and larval mosquito density on the parasite success has been tested through two different experiments. In a first experiment, the density of adults releasing oocysts in water habitats was controlled. They were allowed to released oocysts through feces and body of dead individuals for a week before rearing larvae. In a second experiment, we tested density dependent prophylaxis. To that end, larvae were infected with the same water for 2 days before being separated in batches of 20, 50 and 100 individuals in 100ml of water. Each experiment was repeated five times. The parasite abundance was estimated on emerging adults by qPCR. (B) The impact of larval density (*i.e.* number of larvae per 100ml of water) and (C) the impact of adults density that released oocysts in the larval water habitat on the parasite abundance was represented for each repeat (R1 to R5) and both sexes. Each dot represents a single mosquito. The p-value obtained from a Wald χ^2^ test conducted on a GLMM model represents the significance of the slope.

### The parasite is not likely to impair mosquito dispersion through flight since parasitized individuals are more active

The flight behavior of uninfected and infected mosquitoes was assessed using a Locomotor Activity Monitor. We recorded individually the number of movements of parasitized and unparasitized mosquitoes during 24h (**Figure 6A**). Contrarily to our expectations, parasitized mosquitoes were significantly more active than unparasitized ones (emmeans estimates: 1.34±0.45 and 0.89±0.45 for females and males respectively; **Figure 6B**, **Table 1**). Although it was not statistically significant, the same pattern was observed for parasitized males (49.4 moves) in comparison to unparasitized ones (18.9 moves). Therefore, those results suggest that the low infection rate of mosquitoes in young populations is not related with a higher propensity of uninfected individuals to disperse by flight but the parasite could, conversely, promote the mosquito flight activity.

**Fig. 6.**
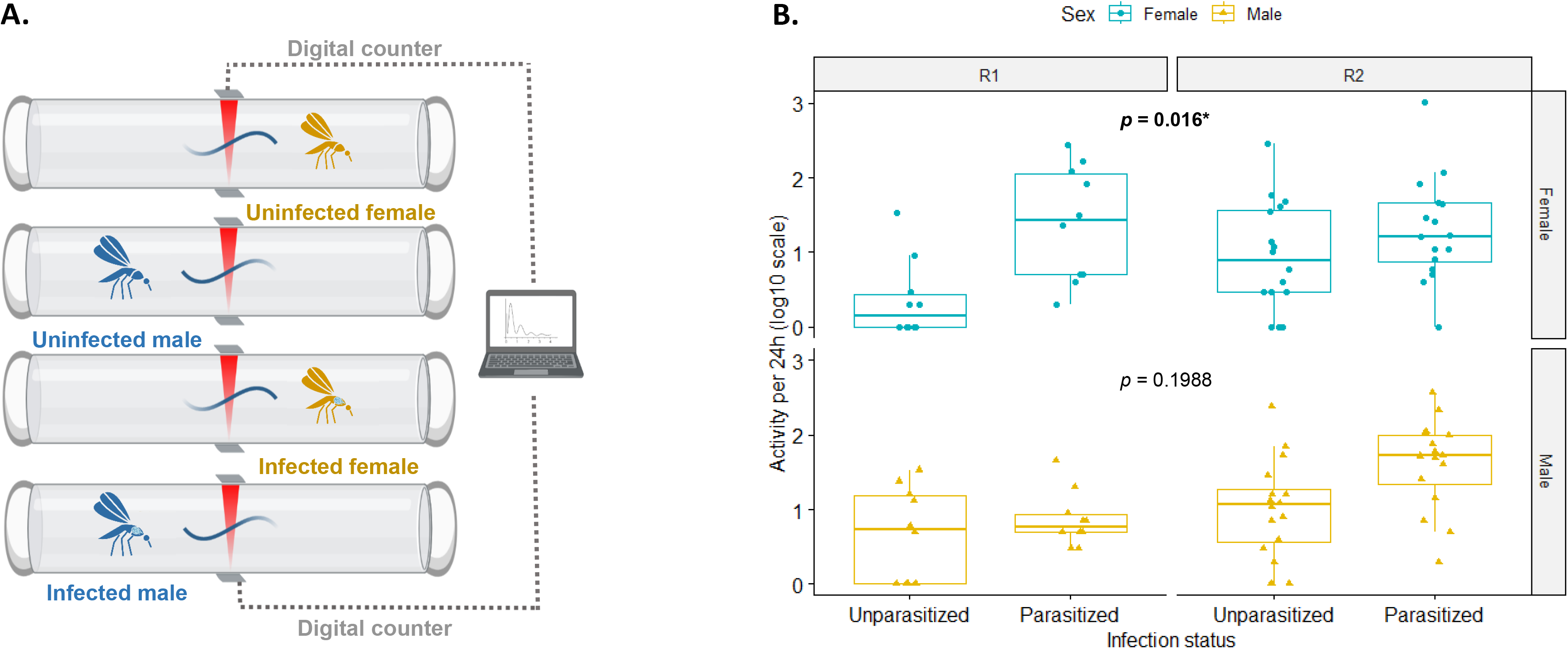
*Ascogregarina taiwanensis* infection impacts on mosquito flight activity. (A) Parasitized or unparasitized mosquitoes activity was measured in a Locomotor Activity Monitor. Movements were recorded for 24h by measuring the number of times each individual pass through an infrared beam. The experiment was repeated on two generations. (B) The impact of parasitism on flight activity (*i.e.* number of time they passed in front of an infrared beam during 24h) was measured for two generations (R1 and R2) and both sexes. The p-value obtained from a Wald χ^2^ test conducted on a GLMM model represents the significance of the slope.

## Discussion

In this study, we reported a positive correlation between *As. taiwanensis* prevalence and *Ae. albopictus* population age for different mosquito populations. Such parasitism escape has been evidenced in particular within recently introduced populations in France and USA. The same pattern was observed regarding the abundance of 2 out of the 3 main *As. taiwanensis* genotypes identified in the metabarcoding analysis. All in all, those convergent tendencies suggest that *Ae. albopictus* might have escaped its natural parasite multiple time along its invasion history. Such decrease might result from subsampling of non-or poorly-infected individuals in the source population that formed the introduced ones. However, this is more likely to occur if the parasite is rare in the source population (16,17) while our field study suggest that the native or old populations sampled along potential source areas are highly colonized by the parasite. Indeed, most recent mosquito records and population genetic studies suggest that mosquito introduction in mainland France is the combination of south European, north-east American introduced populations as well as old invasive populations from the Indian Ocean and native populations from Asia (74,75). Based on PCR diagnostics and microbial community analysis on populations from those areas, we can argue that the parasite is likely to be prevalent and dominant in source population. Three dominant genotypes of *As. taiwanensis* were observed in those area and two of them (Otu0002 and Otu0003) correlated with the laps time separating the mosquito introduction to its sampling. Indeed, metabarcoding and qPCR analysis suggest that those two OTUs dominate the microbiota of intermediate and native populations but not that of recently introduced ones. A recent study on parasite infection in invasive species has shown that parasite characteristics is reliable to predict parasite retention (76). The authors even demonstrated that specialist parasites (like *As. taiwanensis* is) are usually more persistent to co-invade with their native host. All in all, we cannot exclude that stochasticity due to subsampling of non-infected individuals may have played a role in the absence of infection in recently introduced populations but since it is unlikely, we argue that other mechanisms have probably led them to escape *As. taiwanensis*.

The Asian tiger mosquito invasive process is mainly induced by human-mediated large scale dispersal of desiccated eggs as previously discussed (77) and is therefore facilitated by desiccated egg ability to resist without hatching during transportation (78,24). The desiccation and storage laps time that we selected (*i.e.* up to 1month) were included within the range the eggs can survive without showing consistent decrease in survival (40). We demonstrated that the abundance and prevalence of parasite infection in mosquitoes hatched from desiccated eggs decreases as a function of the time the eggs were kept before being flooded. Similarly, the gregarine *Apolocystis elongata* showed a reduced infectivity toward its earthworm host *Eisenia foetida* three weeks after oocysts have been released on a dry soil (79). Sensitivity to desiccation has also been reported as a major oocyst weakness in related Apicomplexan species such as *Cryptosporidium* or *Eimeria* species (80,81). The molecular mechanism involved in desiccation sensitivity has poorly been studied until now in Apicomplexa but is likely to be due to the peculiar evolution of those microorganisms that recently diverged from marine algae (82). At the opposite, resistance of cysts to desiccation have been acquired in some Apicomplexan taxa such as *Toxoplasma* spp. and was associated to the production of Late Embryogenesis Abundant (LEA)-related proteins (83). The particular biochemical properties of those protein enable them to harbor chaperone-like activities and contribute to protein stabilization that may reinforce the cyst wall integrity and to limit water loss. The lack of acquisition of such protein in *As. taiwanensis* still needs to be demonstrated but may explain why this parasite that is often released in water container poorly resist to desiccation.

Since transportation of desiccated eggs is a major path driving the mosquito introduction (29–32,74,84,85), human-aided transportation could be a consistent factor explaining *Ae. albopictus* parasitism escape after mosquito population introduction. However, part of the mosquitoes are also transported as adults for a shorter period of time in car cabin and this path that can hardly be related to parasitism escape (29). Habitat and host permissiveness to the parasite after introduction may also be a key factor that could have decreased the infection probability for a short period of time after mosquito introduction (86). Those processes are complex and may involve biotic and abiotic components. We did not fully address this question and argue that further studies must be conducted to better understand the importance of genotype x genotype x environment (GxGxE) interactions in *Ae. Albopictus* - *As. taiwanensis* interactions. However, our results provide new insights for further investigations. Firstly, the metabarcoding of eukaryotic and prokaryotic communities in *Ae. albopictus* did not evidence that the parasite was replaced by another competitive parasite from the invasion range. This part suggest that the biotic part associated to microorganism does not seem very relevant to explain the host-parasite dynamics. Secondly, the habitats in mainland France and in USA could be partially refractory to the parasite since we evidenced for both localizations’ correlation between geographical distances and parasite prevalence. Further studies on biotic and abiotics are necessary to conclude on the environment importance on this host-parasite relationship.

In other host-parasite systems such as snail-trematode, the host density has been shown to be a good predictor of parasite colonization success (87). Such phenomena are often predicted to occur for horizontally transmitted parasites due to inter-individual contacts leading to higher transmission probability that increase with population density (88). Since inter-individual horizontal transmission of parasites is mediated through oocysts released from adults in water habitats, one can expect that a reduced number of mosquitoes invading a new area would result in a limited number of parasites in the breeding sites to colonize new hosts. Previous studies on *As. taiwanensis* evidenced that the parasite oocyst load released in water influence the success of host infection (59). In this study, we have complemented those results by showing that the number of adult mosquitoes releasing parasites positively increases the number of parasites efficiently colonizing unparasitized conspecifics emerging from water habitats. Interestingly an inter-seasonal mosquito and *As. taiwanensis* dynamics have been shown to be correlated over the year (51). Those results converge toward the importance of high mosquito population densities for an efficient parasite transmission, independently from the probability of parasite acquisition. Previous studies on gregarious insects showed that an adaptive response to host high density leads to a reduction of the parasite susceptibility through the enhancement of their immune system (22). This phenomenon is also named Density-Dependent Prophylaxis (DDP). For instance, in response to density raise, the bumble bee *Bombus terrestris* (Hymenoptera) and the velvet bean caterpillar *Anticarsia gemmatalis*, increase their phenoloxidase activity (immune response leading to a melanization process) (89,90). In this study, we have shown an absence of larval density-mediated modulation of the parasite infection cycle. Although the immune response has not been tested, we can then predict that the parasite transmission drastically increases with the population size without being slowed down by DDP.

We have highlighted the impact of the different invasion processes in the host-parasite relationship. Contrarily to our hypothesis, local mosquito dispersal through flight has been shown to be improved by parasitism. Local flight distance of field populations is generally limited to a few hundred meters (47) and remains a secondary factor to explain *Ae. albopictus* invasiveness. The hypothesis of gregarine dispersal through local flight is debatable due to opposite examples. Indeed, the infectivity of the gregarine *Ophryocystis elektroscirrha* to monarch butterfly (*Danaus plexippus*) has been proved to reduce host wing mass and resistance (91). In the dragonfly (*Libellula pulchella*), gregarines are also negatively affecting the flight performance by decreasing the concentration of muscle myosin and impairing muscle contractile performance (92,93). The results reported here are preliminary. Indeed, laboratory experiments on mosquitoes behavior using locomotor activity monitor do not take into account various stimuli (e.g. breeding sites, hosts…) from the environment that might differentially enhance the local dispersal of mosquitoes (94,95). Other factors such as the mosquito age and nutrition would also be considered in the future while previous studies showed that the activity varies according to those factors (96). If our result is confirmed under more complex conditions, this would suggest that mosquitoes that have been colonized by the parasite after introduction might be more prompt to locally disperse.

Recently, a mathematical model predicted *Ae. albopictus* competitiveness against the tree hole mosquito *Ae. triseriatus* in presence of gregarine parasites (97). In most scenarii, *Ae. albopictus* dominates *Ae. triseriatus*, but *As. taiwanensis* infection may decrease the competitiveness of its host by increasing the larval development time and inducing a higher mortality, according to other studies (98). The authors concluded that parasitism reduces *Ae. albopictus* competitiveness. Selective pressure induced by competition might explain the loss of parasites in recently introduced *Ae. albopictus* populations since its presence might restrain the population settlement probability. Another hypothesis to explain the parasitism escape is a dilution effect when *Ae. albopictus* colonizes new areas. Both parasite prevalence and intensity could be reduced due to a wider range of potential host (induced by native mosquito species). Indeed, they can colonize first instar larvae of other mosquito species but rarely achieve to complete their life cycle and replicate inside them (49). To the best of our knowledge, no study evidenced dilution of *As. taiwanensis* out of *Ae. albopictus*. However, dilution effect of *Ascogregarina barreti* has been experimentally observed during co-inoculation of its native host *Ae. triseriatus* with other mosquito species such as *Ae. japonicus* (99) and *Ae. albopictus* (100). It was also suggested by field surveys reporting that the prevalence of *As. taiwanensis* colonizing *Ae. albopictus* was higher when the mosquitoes did not share breeding sites with *Ae. aegypti* (101)*. Ae albopictus* is often reported as a strong competitor toward other mosquitoes (102–107). Therefore, the parasite prevalence dynamics in young populations could be due to an evolution of the dilution effect resulting from a higher diversity of mosquitoes in breeding sites at the early stage of the colonization process that decrease over the time due to interspecific competition. Field studies on the arthropod’s community sharing breeding sites with recent and ancient populations may help to find relevant alternative hosts to test this hypothesis.

## Conclusion

In this study, we provide new insights into the dynamics of *Ae. albopictus*-*As. taiwanensis* relationship by combining field observations and laboratory experiments. In French populations, the mosquito host has escaped its parasite after introduction and this process is unlikely to be stochastic. The maintenance of non-flooded eggs in containers during human-aided transportation and the decrease in mosquito founding population after introduction have likely altered the parasite maintenance and transmission. Local dispersal through flight is probably not affected by the parasite since the flight activity of unparasitized individuals was lower than that of their parasitized conspecifics. We need further investigations on the consequences of the rupture of this natural interaction to understand how it may affect the mosquito ecology, population dynamics and pathogen transmission.

## Acknowledgement

Part of the mosquito collection was enabled by the international network of the ‘Mosquito Microbiome Consortium’ (https://mosquito-microbiome.org/). This project was funded by the French National Research Program for Environmental and Occupational Health of Anses, grant number 19-95. We would also like to thank the EC2CO program for its support to this project (project name ‘INTERASCO’) as well as the IDEX Breakthrough (project name ‘MICRO-BEHAVE’). Sampling collections were also supported by the Thailand Program Management Unit for Human Resources & Institutional Development, Research and Innovation (PMU-B), NXPO, grant number B17F640002. This study used resources from the iBio platform and the FR BioEEnViS. Finally, the authors would like to thank James Swanson and Stephanie Jiang from Emory University for their help during sampling.

## Data availability

All the reads, datasets and scripts used for this analysis have been deposited on zenodo (https://zenodo.org/record/8252320).

## Supplementary material

### DNA extraction

Mosquito samples were rinsed three time with sterile water, surface disinfected 5min in ethanol and washed 5 times in sterile water. DNA was extracted from each mosquito individually crushed with three 3mm borosilicate beads and under agitation during 60s at 6m/s with a Fastprep homogenizer (MP Biomedicals). Homogenate was then suspended into a CTAB extraction buffer (2% hexadecyltrimethyl ammonium bromide, 1.4 M NaCl, 0.02 M EDTA, 0.1 M Tris pH 8.0, 0.2% 2-β-mercaptoethanol) and heated for 2h at 60°C. After incubation, 2mg/ml of RNase was added to the extraction buffer and incubated at 37°C for 5 min. DNA was separated from proteins and lipids with an equivalent volume of phenol-chloroform-isoamyl alcohol (25:24:1, v:v:v), vortexed for 15min and centrifuged for 30min at 16,100g. Aqueous phase was harvested and mixed with an equivalent volume of chloroform:isoamyl alcohol (24:1, v:v). The mixture was vortexed for 15min and centrifuged for 30min. The newly separated aqueous phase containing the nucleic acids was harvested. DNA was precipitated with isopropyl alcohol and centrifuged for 30min at 16,100g. DNA pellet were washed with 70% cold ethanol and centrifuged for 30min at 16,100g. Ethanol was removed and DNA was suspended in 20µl of sterile water. DNA extracts were quantified with a spectrophotometer (SAFAS) and diluted at 15ng/µl.

### qPCR estimation of the eukaryotic and prokaryotic microbial density

Since variations in OTU abundances may be influenced by variations in absolute densities of microorganisms, qPCR essays were performed on samples for which variations in *As. taiwanensis* abundances were observed. Following the metabarcoding analysis and correlation network analysis, three different groups of *Ae. albopictus* microbiota were observed depending on their infection status with *As. taiwanensis* (see Results section). Prokaryotic and Eukaryotic qPCR essays were performed on 7 males and 7 females from each of those three groups using the same primers than in metabarcoding. Reaction mix contained 1X of SYBR Green iTaq (Bio-Rad), 3µM of each primer and 5ng of DNA. PCR cycles were performed on a CFX96 Touch Real-Time PCR detection system (Bio-Rad) following a 3min initial denaturation at 95 °C followed by 40cycles with 5s of denaturation at 95°C and 1min of hybridization and amplification at 60°C. A final denaturation of the PCR products was conducted by progressively increasing the temperature from 60°C to 90°C at 0.11°C/s. The Ct values were used to quantify the copies number of each gene with a standard range of 10^7^ to 10^0^ genomic DNA copies of *Escherichia coli* DH5-α for the 16S rDNA and a 10^7^ to 10^0^ genomic DNA copies of *Saccharomyces cerevisiae* CEN.PK2-2 for the 18SrDNA.

### PCR-based estimation of Ascogregarina taiwanensis prevalence

Detection of *As. taiwanensis* in DNA from field collected individuals was conducted with diagnostic PCR using primers targeting the ITS1-5.8SrDNA-ITS2 intergenic region with the previously published AU (5’-ACCGCCCGTCCGTTCAATCG-3’) and AT (5’-GAGAAGCCGTCGTCAATACAGC-3’) primers (Morales et al., 2005). The mix contained 1X of Hoststart Master mix (Dutscher, 0.2ng/ml of BSA (Promega), 0.1µM of each primer, 0.1X of GC-Enhancer (Biofidal) and 30ng of mosquito genomic DNA. Reactions were performed with a 10min initial denaturation at 96°C, followed by 35 cycles of amplification including 20s at 96°C, 1min at 50°C, 2min at 72°C and a final elongation of 10min at 72°C. Amplification signals were identified under UV light on a Geldoc 2000 system (Biorad) after 20min of electrophoresis migration at 100V on a 1.5% agarose gel colored with Ethidium Bromide.

### Design of qPCR primers for the quantification of Ascogregarina taiwanensis

Primers hybridizing *As. taiwanensis* were designed *in silico* based on an alignment of previously published 18S rDNA sequences of Apicomplexa (genebank accession numbers

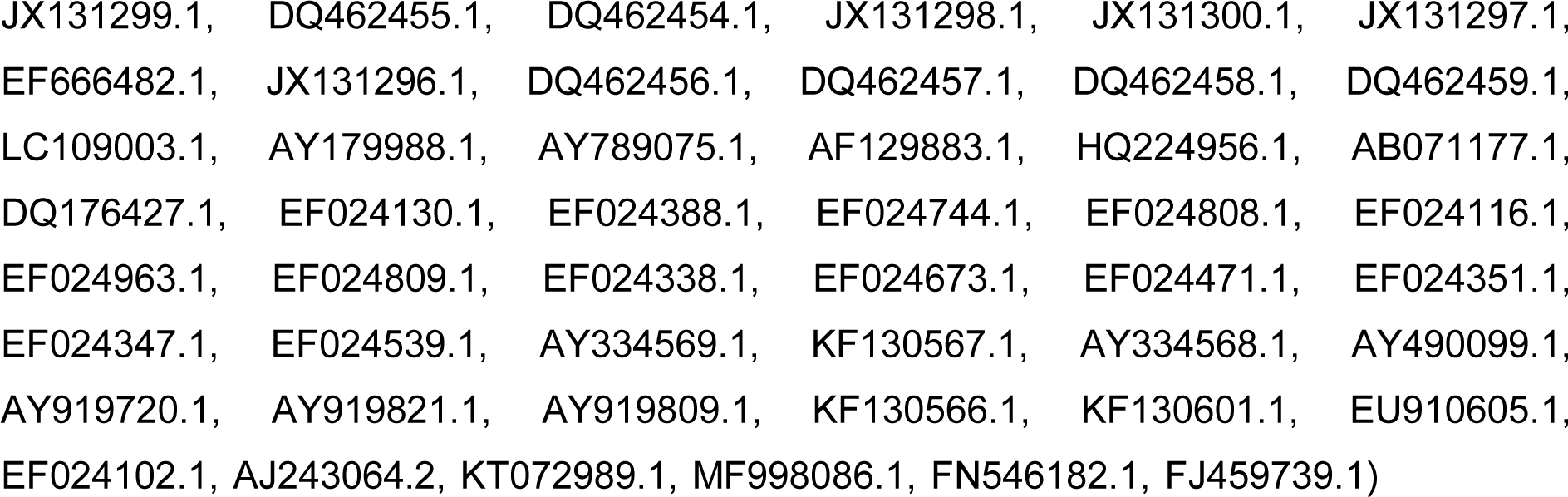

to select a conserved region in *As. taiwanensis* that diverged from other species. Two specific primers named AscoJSF (5’-CTMGGCTTGACTTCGGTCT-3’) and AscoJSR (5’-TTCCATGCTGGAGTATTCAAGG-3’) were designed and produced a fragment of 150bp. A standard concentration of the fragment cloned in a vector was prepared. It was first amplified with a PCR mix containing 1X of buffer, 1.5mM of MgCl_2_, 200µM of dNTP, 200nM of each primer, 0.625 Units of Taq DNA Polymerase (New England Biolabs) and 30ng of DNA from a parasitized mosquito individual. The PCR program was ran with 95°C of initial denaturation for 4min, followed by 35 cycles including 30s at 95°C, 30s at 56°C, 15s at 68°C and a final elongation of 5min at 68°C. The PCR product was purified with a PCR purification kit MinElute (Qiagen), cloned into a PCR 2.1-TOPO TA vector (Invitrogen) and transferred into competent *Escherichia coli* Top10 cells (Invitrogen) following manufacturers’ recommendations. Competent cells containing the vector and the insert were selected on LB agar supplemented with 25µg/ml of Kanamycin and 20µg/ml of Xgal and 23.8µg/ml of IPTG after 24h of incubation at 37°C. Presence of the insert was also verified with PCR on white clones that developed on the medium. DNA from the competent cells was extracted with a DNeasy Blood and Tissue kit (Qiagen) following manufacturers’ recommendations and linearized 1h at 37°C in a mix containing 0.2µg/µl of DNA, 1X of SurE/Cut Buffer B and 10U of BamH1 endonuclease (Roche). Reaction was inactivated at 65°C for 15min. This digested DNA was purified with a MinElute PCR purification kit (Qiagen), quantified with Quanti-it dsDNA BR Assay kit (Invitrogen) and used as a standard for qPCR with concentrations ranging from 10^8^ to 10^0^ copies. Reaction mixes for quantification included 0.5µM of each primer, 1X of iTaq Universal SYBR Green Supermix (Bio-Rad) and 5ng of DNA. The qPCR program included a 5min initial denaturation at 95°C, followed by 40 cycles with 15s at 95°C, 30s at 56°C and 10s at 60°C followed by a progressive denaturation of the qPCR products from 65°C to 97°C at 0.11°C/s.

### Metabarcoding of eukaryotic and prokaryotic microbial communities

Both PCR amplifications were conducted under sterile conditions with 1X of Hotstart 5X Bioamp Master Mix (Biofidal), 0.2µM of each primers, 0.2mg/ml of Bovine Serum Albumin (New England Biolabs), 0.4X of GC rich Enhancer (Biofidal). For 16S rDNA, PCR program started with 10min of initial denaturation at 96°C followed by 35 cycles including 20s of denaturation at 96°C, 1min of annealing at 54°C, 30s of extension at 72°C and ended with a final extension of 10min at 72°C. For 18S rDNA, PCR program started with 10min of initial denaturation at 96°C followed by 35 cycles including 20s of denaturation at 96°C, 30s of annealing at 54°C, 30s of extension at 72°C and ended with a final extension of 10min at 72°C. Three negative controls corresponding to blank extractions and PCR were included for each gene.

Details concerning the mother pipelines used for bioinformatics treatment: A total of 12,836,921 and 14,248,382 reads were obtained from 16S and 18S rDNA respectively and were demultiplexed using the Mothur pipeline (69). Reads were trimmed based on size (270-310bp and 400-550bp for the 16S and 18S rDNA, respectively) and sequences were conserved if they contained no ambiguous position and if both strands of the paired-end reads aligned together. Chimeras were removed using CHIMERAVSEARCH. The alignment was performed using Silva v.132 database for bacteria and eukarya and reads that did not align were filtered out. However, for sequences that matched gregarine, a supplemental local alignment (BLAST) was performed against the Genbank database due to the lack of reference sequences in Silva. Clustering was performed using Opticlust method allowing a maximum dissimilarity rate of 3%.

---

**Figure S1.**
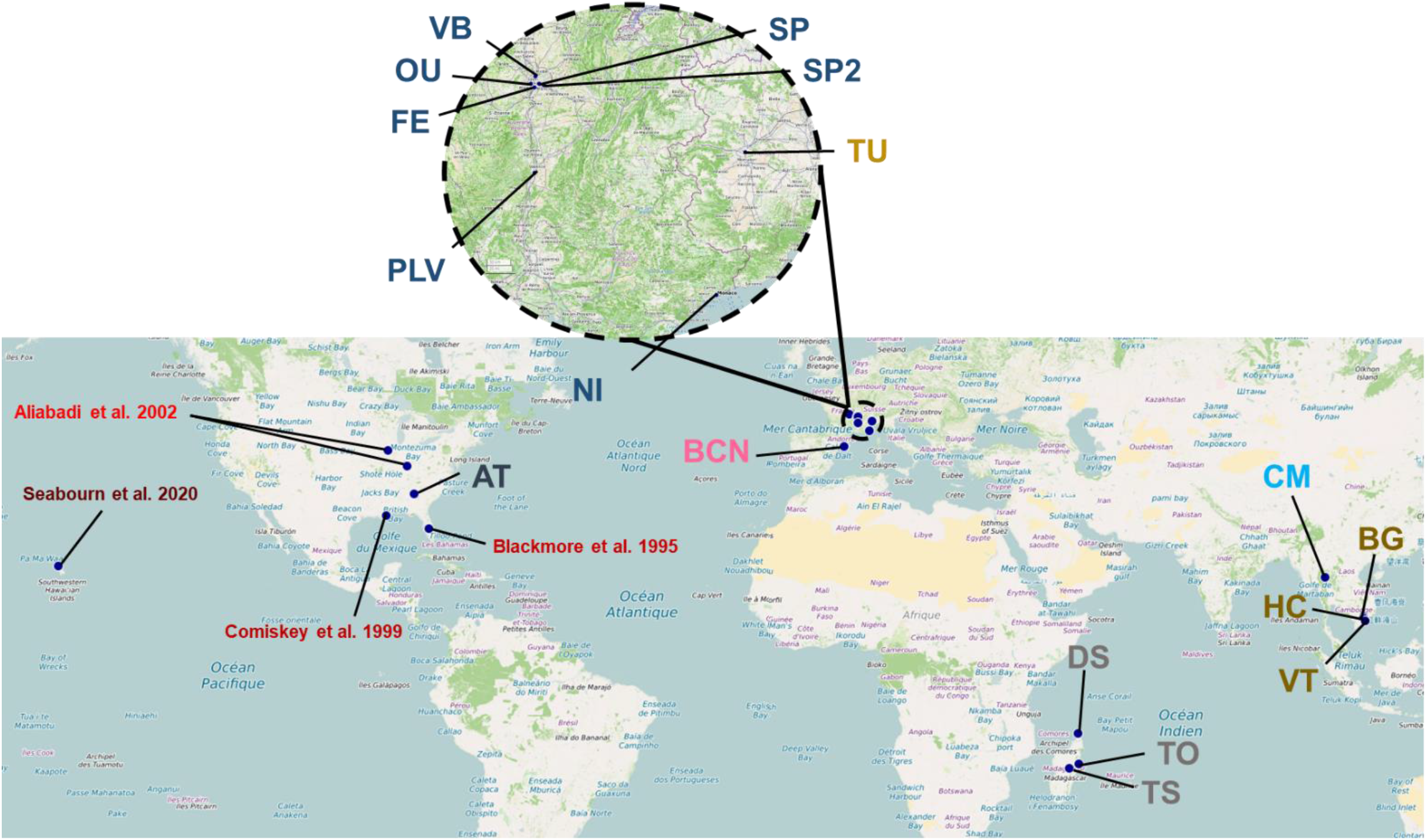
World map representing the collection sites of mosquitoes’ populations used in this study.

**Figure S2.**
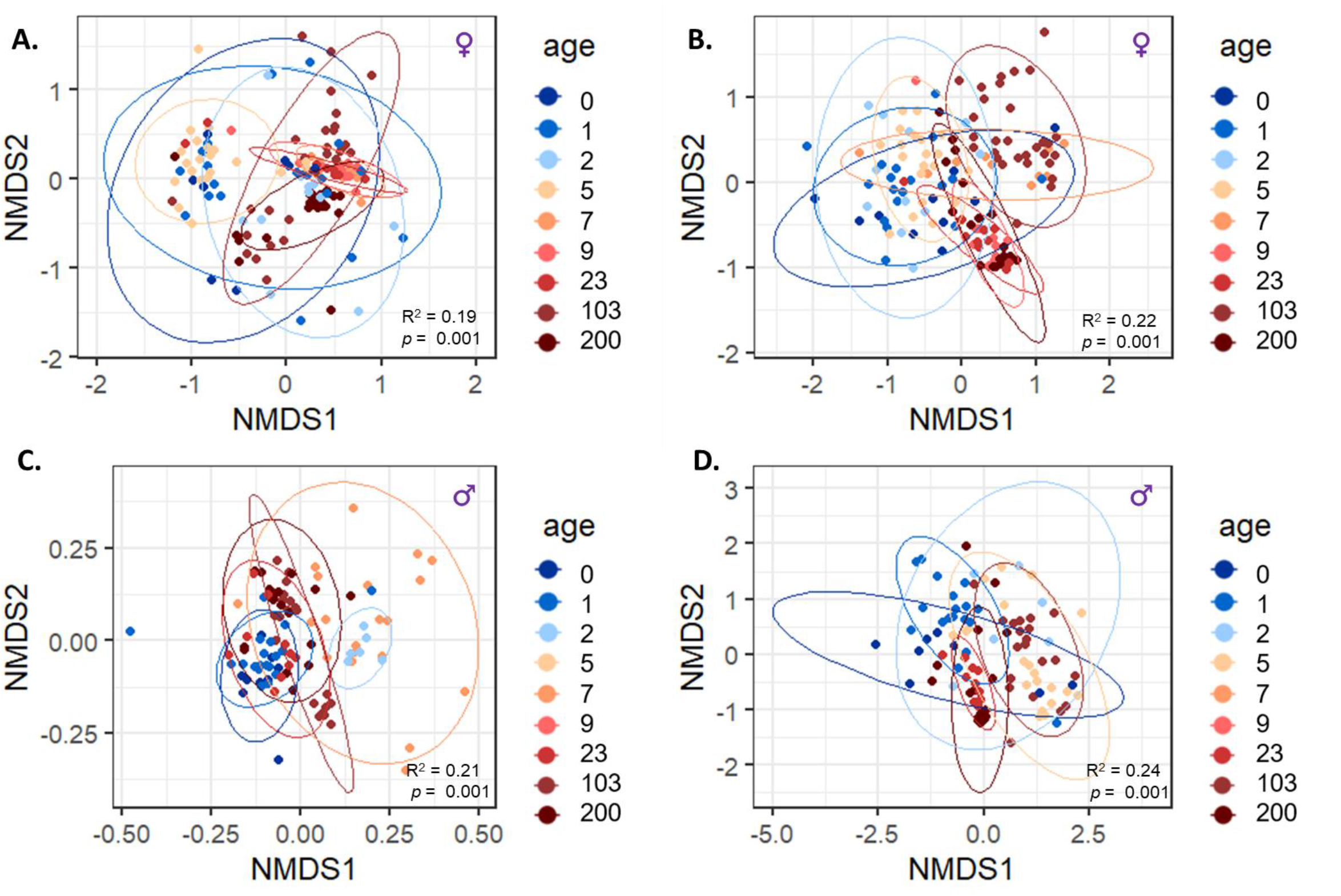
Microbiota composition dissimilarity separation according to mosquito population ages. For both (A, B) female and (C, D) male individuals the dissimilarity (bray-curtis index) of (A, C) prokaryotic and (B, D) eukaryotic microbiota communities was represented. The points are colored with a gradient ranging from blue to red based on the age of each population (years after introduction). Native populations were set to 200 years.

**Figure S3.**
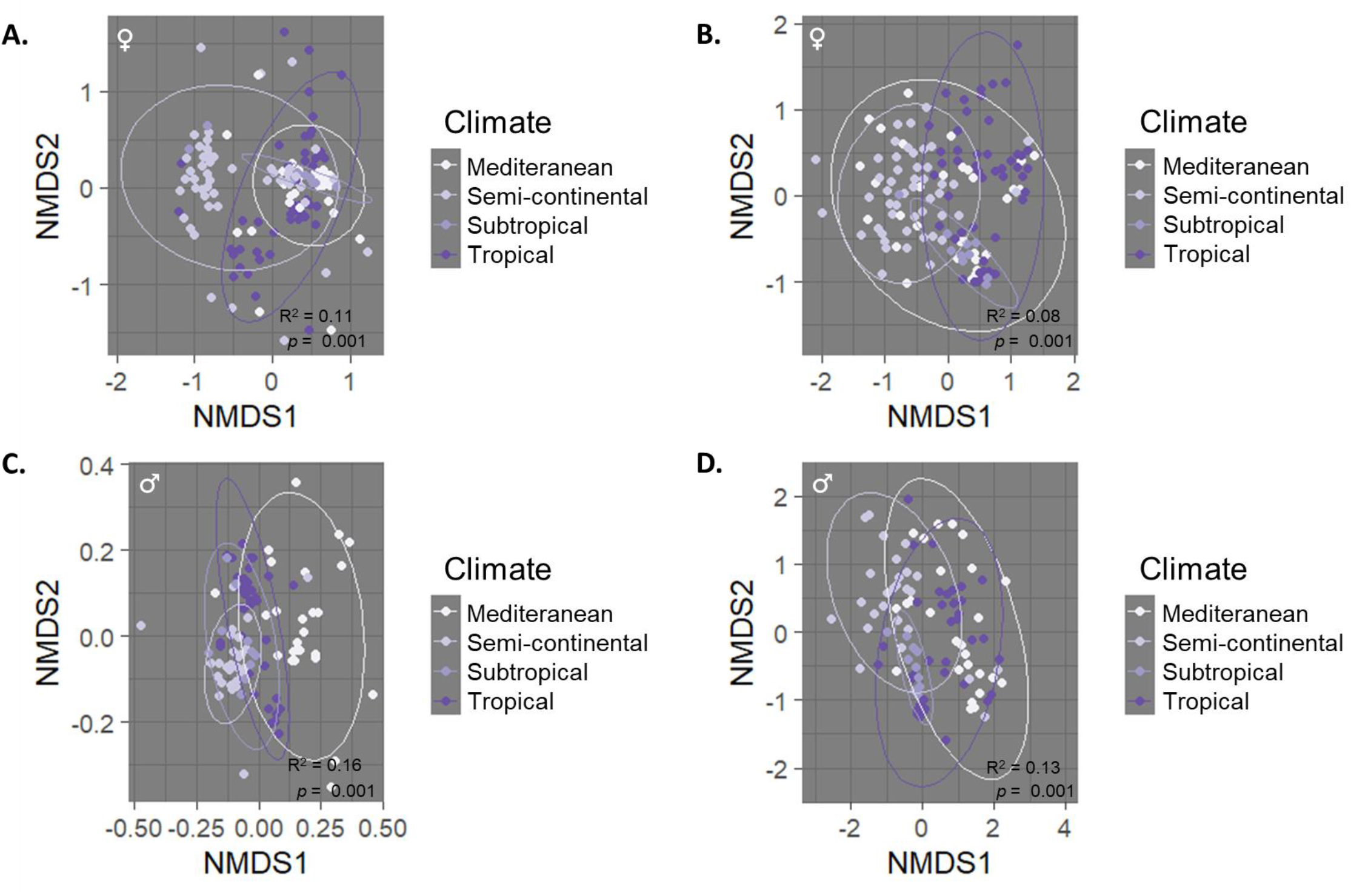
Microbiota dissimilarity separation according to climate. For both (A, B) female and (C, D) male individuals the dissimilarity (bray-curtis index) of (A, C) prokaryotic and (B, D) eukaryotic microbiota communities was represented. The points are colored with a gradient ranging from blue to red based on climatic regions. Native populations were set to 200 years.

**Figure S4.**
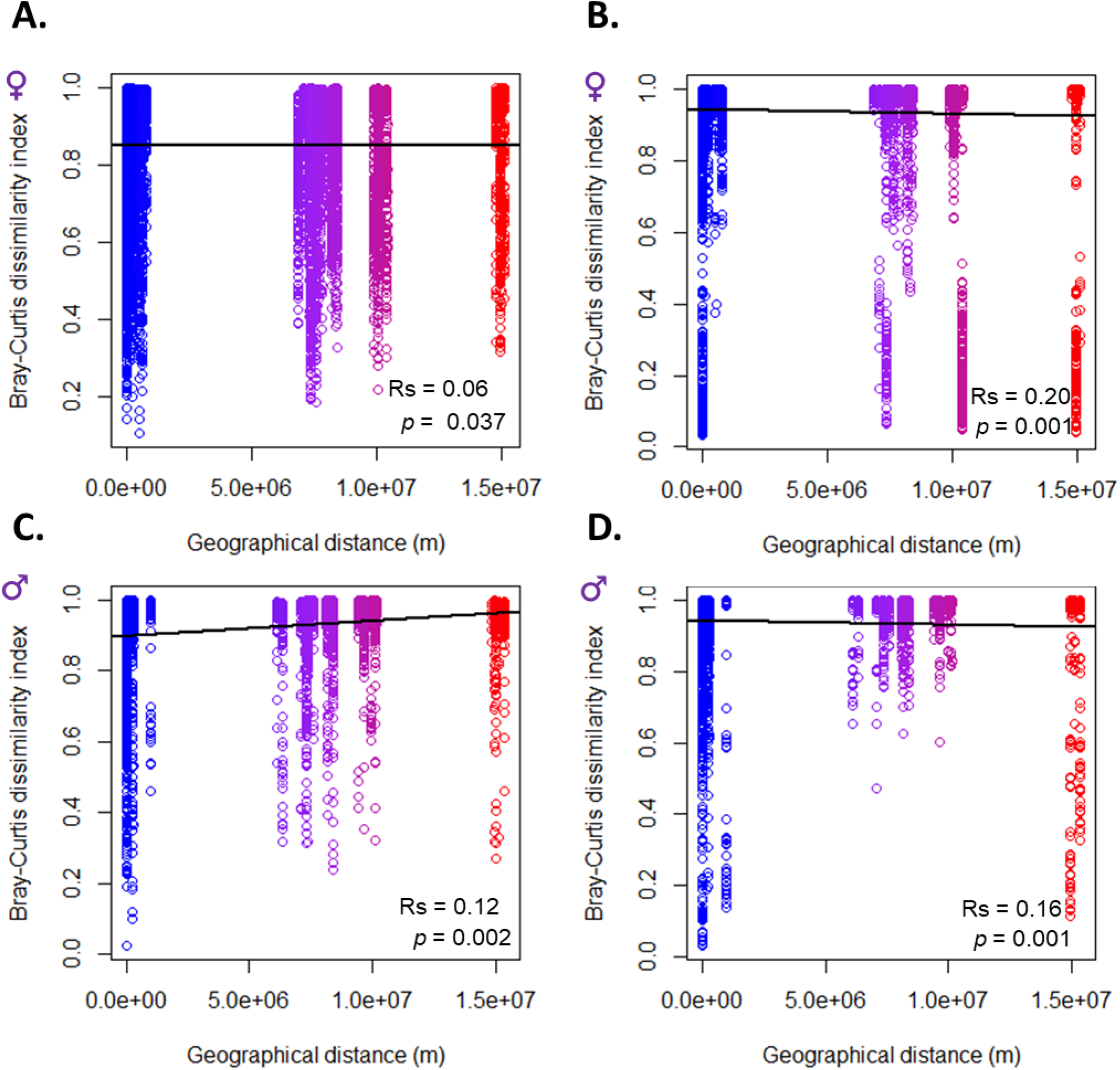
Microbiota dissimilarity correlation with geographical distances. Spearman correlation analysis between the microbiota community dissimilarity estimated with the Bray-Curtis index and geographical ellipsoid distance. Correlations were represented for both (A, B) female and (C, D) male individuals and for (A, C) prokaryotic and (B, D) eukaryotic microbiota communities. The points are colored with a gradient ranging from blue to red based on geographical distance between both points being compared. The correlation coefficient (Rs) and p-value are represented on each graph.

**Figure S5.**
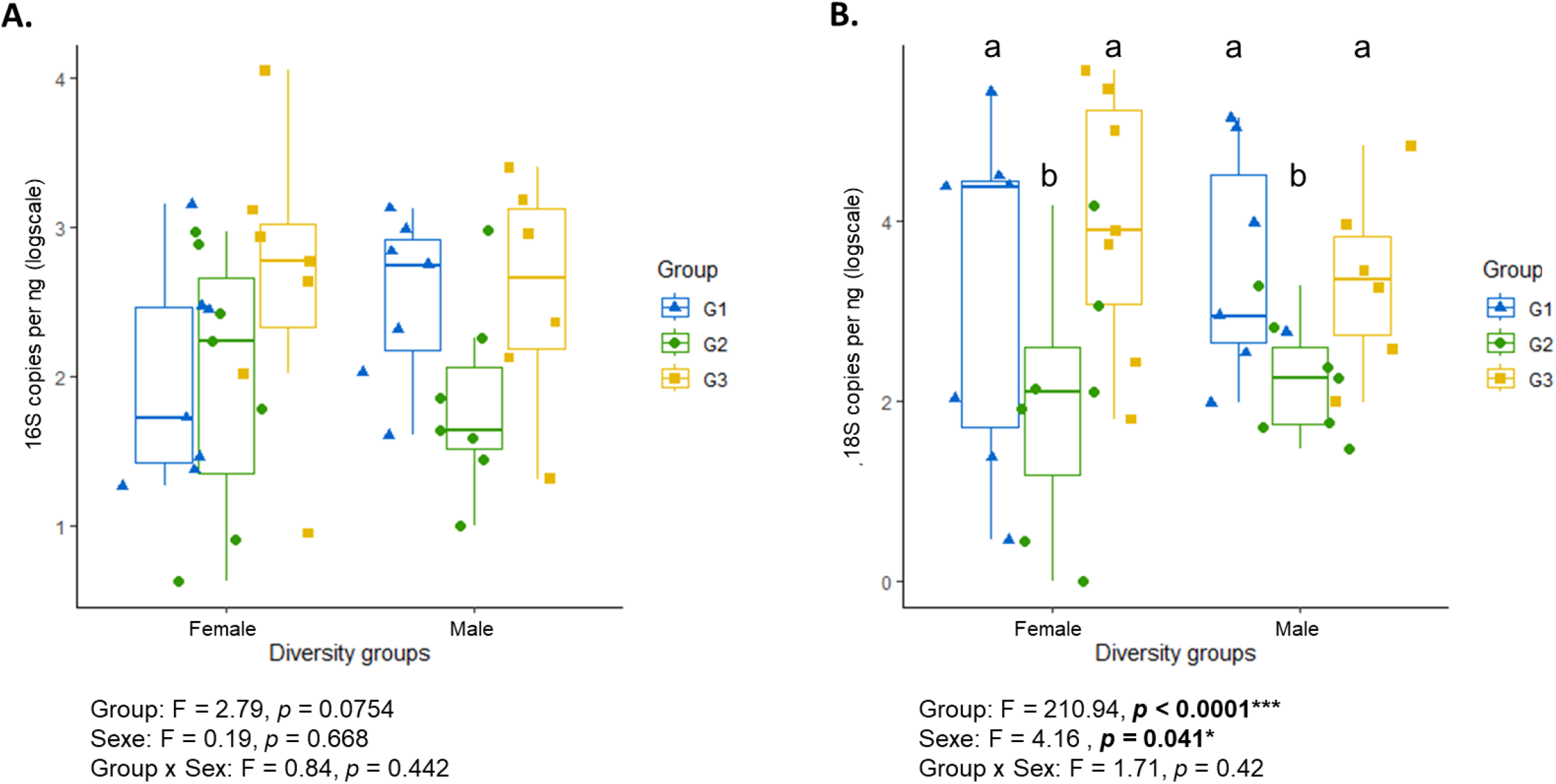
Quantification of microbial densities. Quantifications were performed through qPCR of the (A) 16S rDNA genes for the eukaryotic community or (B) 18S rDNA gene for the prokaryotic community. Females and males are respectively colored in red and blue. G1 represent samples from various origins for which the *Ascogregarina taiwanensis* Otu000001 dominates the microbiota. G2 represents samples for which *As. taiwanensis* Otu000002 and Otu000003 dominates the microbiota. Finally, G3 represents samples that are not dominated by *As. taiwanensis*.

**Table S1.**
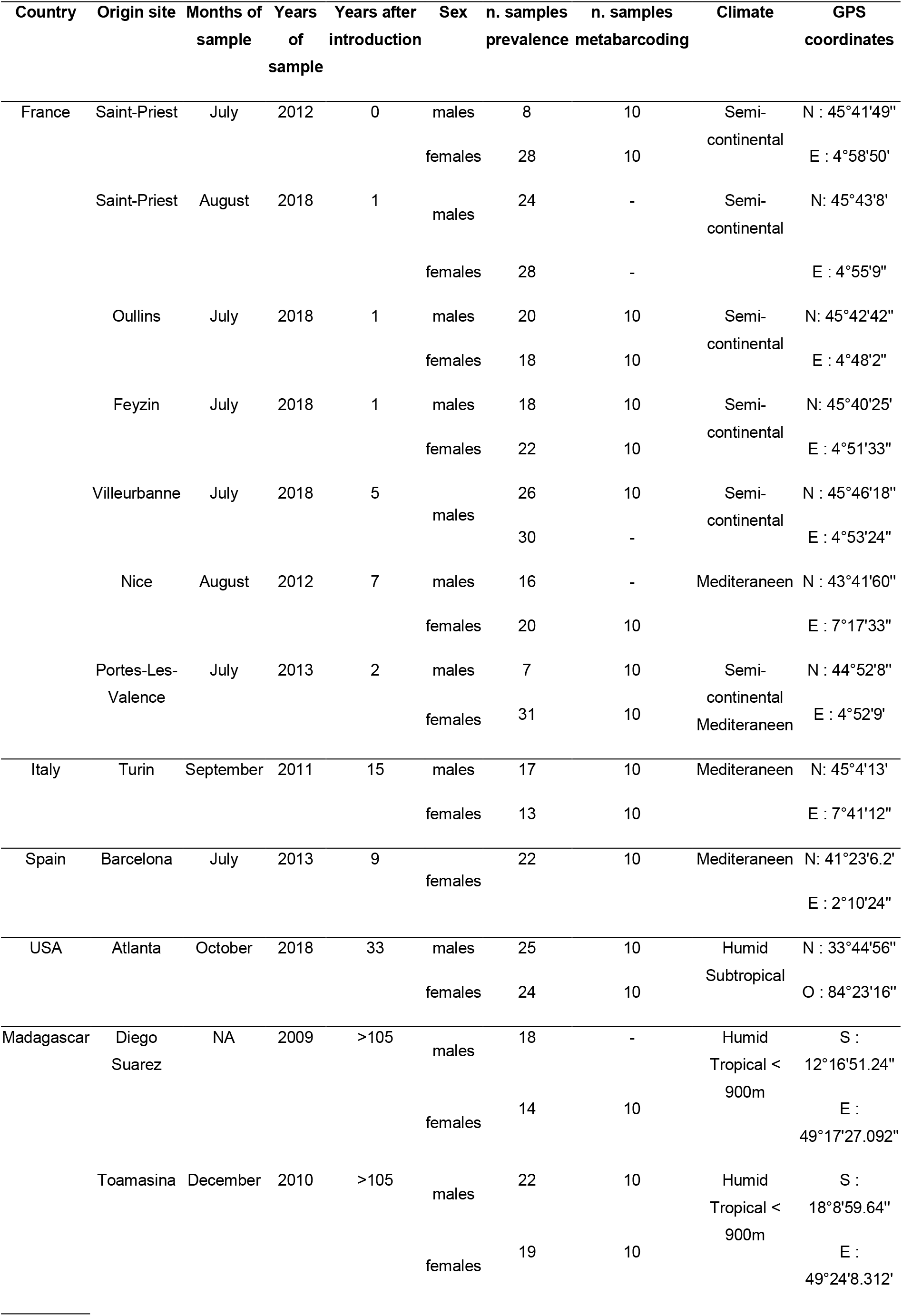
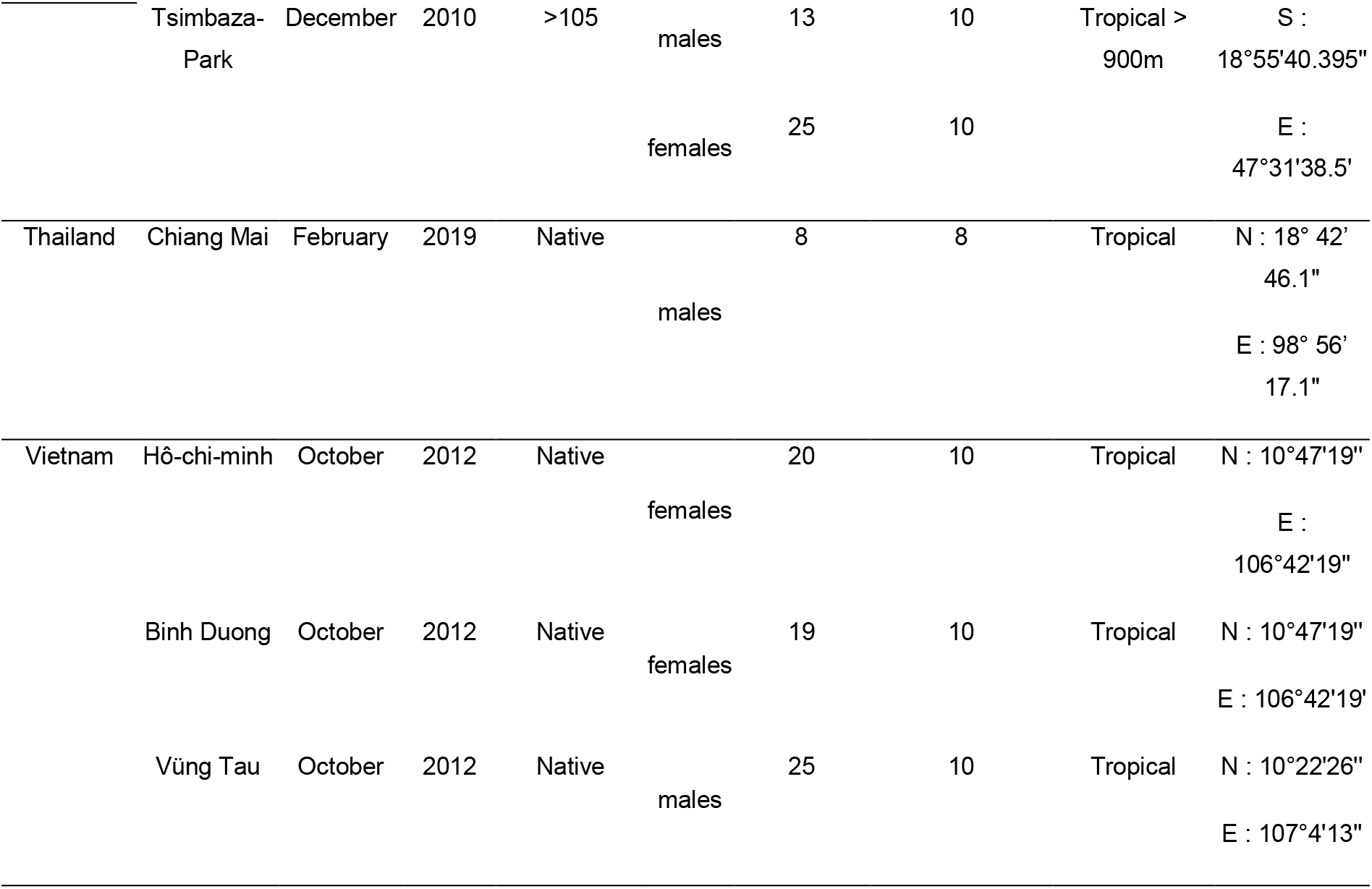
Aedes albopictus population used in the current study.

| Country | Origin site | Months of sample | Years of sample | Years after introduction | Sex | n. samples prevalence | n. samples metabarcoding | Climate | GPS coordinates |
| --- | --- | --- | --- | --- | --- | --- | --- | --- | --- |
| France | Saint-Priest | July | 2012 | 0 | males | 8 | 10 | Semi-continental | N : 45°41'49" |
|  |  |  |  |  | females | 28 | 10 |  | E : 4°58'50' |
|  | Saint-Priest | August | 2018 | 1 | males | 24 | - | Semi-continental | N: 45°43'8' |
|  |  |  |  |  | females | 28 | - |  | E : 4°55'9" |
|  | Oullins | July | 2018 | 1 | males | 20 | 10 | Semi-continental | N: 45°42'42" |
|  |  |  |  |  | females | 18 | 10 |  | E : 4°48'2" |
|  | Feyzin | July | 2018 | 1 | males | 18 | 10 | Semi-continental | N: 45°40'25' |
|  |  |  |  |  | females | 22 | 10 |  | E : 4°51'33" |
|  | Villeurbanne | July | 2018 | 5 | males | 26 | 10 | Semi-continental | N : 45°46'18" |
|  |  |  |  |  |  | 30 | - |  | E : 4°53'24" |
|  | Nice | August | 2012 | 7 | males | 16 | - | Mediterranean | N : 43°41'60" |
|  |  |  |  |  | females | 20 | 10 |  | E : 7°17'33" |
|  | Portes-Les-Valence | July | 2013 | 2 | males | 7 | 10 | Semi-continental<br>Mediterranean | N : 44°52'8" |
|  |  |  |  |  | females | 31 | 10 |  | E : 4°52'9' |
| Italy | Turin | September | 2011 | 15 | males | 17 | 10 | Mediterranean | N: 45°4'13' |
|  |  |  |  |  | females | 13 | 10 |  | E : 7°41'12" |
| Spain | Barcelona | July | 2013 | 9 | females | 22 | 10 | Mediterranean | N: 41°23'6.2'<br>E : 2°10'24" |
| USA | Atlanta | October | 2018 | 33 | males | 25 | 10 | Humid<br>Subtropical | N : 33°44'56" |
|  |  |  |  |  | females | 24 | 10 |  | O : 84°23'16" |
| Madagascar | Diego Suarez | NA | 2009 | >105 | males | 18 | - | Humid<br>Tropical < 900m | S :<br>12°16'51.24" |
|  |  |  |  |  | females | 14 | 10 |  | E :<br>49°17'27.092" |
|  | Toamasina | December | 2010 | >105 | males | 22 | 10 | Humid<br>Tropical < 900m | S :<br>18°8'59.64" |
|  |  |  |  |  | females | 19 | 10 |  | E :<br>49°24'8.312' |
|  | Tsimbaza-Park | December | 2010 | >105 | males | 13 | 10 | Tropical > 900m | S : 18°55'40.395" |
|  |  |  |  |  | females | 25 | 10 |  | E : 47°31'38.5" |
| Thailand | Chiang Mai | February | 2019 | Native | males | 8 | 8 | Tropical | N : 18° 42' 46.1" |
|  |  |  |  |  |  |  |  |  | E : 98° 56' 17.1" |
| Vietnam | Hô-chi-minh | October | 2012 | Native | females | 20 | 10 | Tropical | N : 10°47'19" |
|  |  |  |  |  |  |  |  |  | E : 106°42'19" |
|  | Binh Duong | October | 2012 | Native | females | 19 | 10 | Tropical | N : 10°47'19" |
|  |  |  |  |  |  |  |  |  | E : 106°42'19" |
|  | Vũng Tau | October | 2012 | Native | males | 25 | 10 | Tropical | N : 10°22'26" |
|  |  |  |  |  |  |  |  |  | E : 107°4'13" |

**Table S2.**
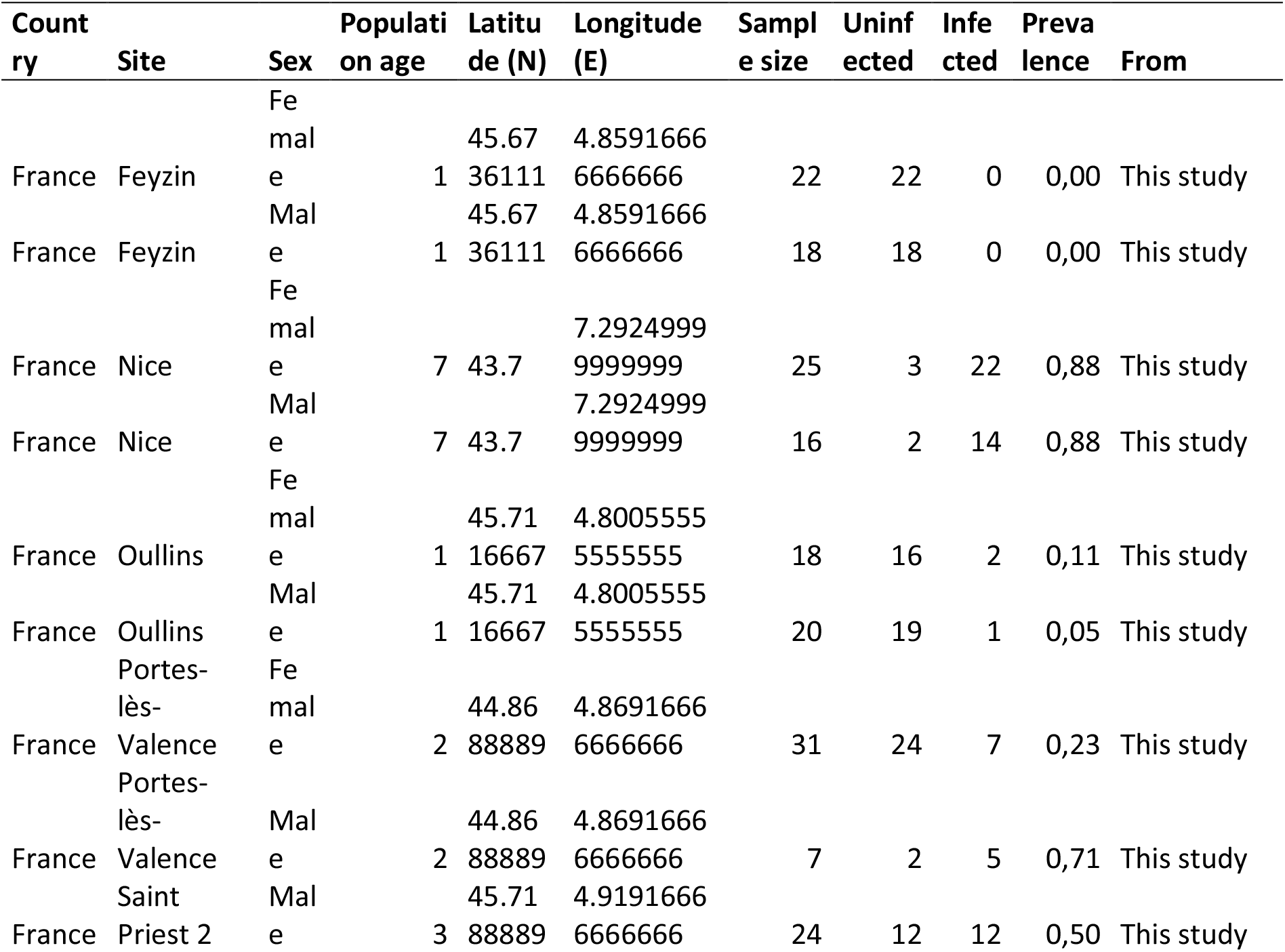
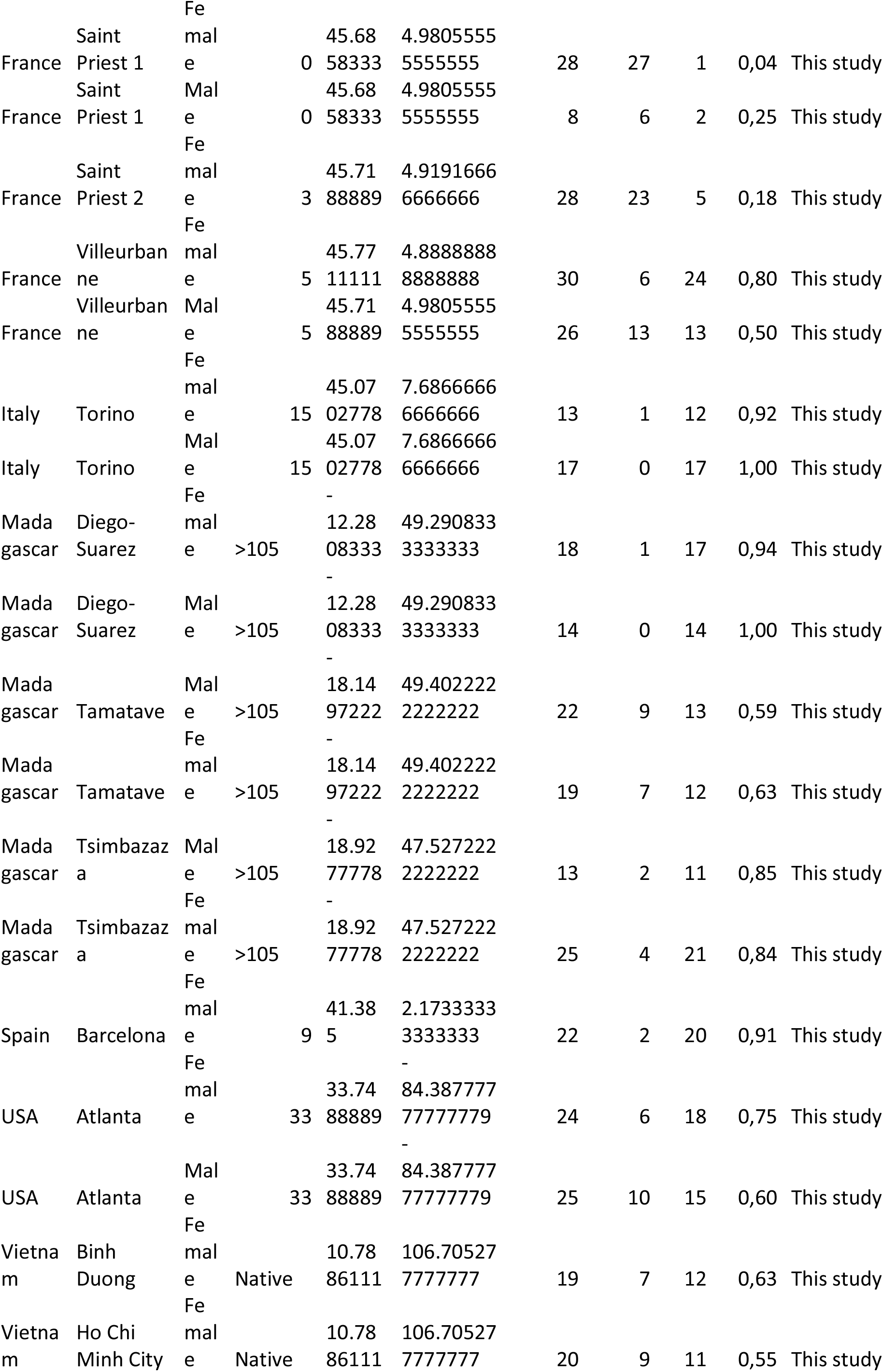
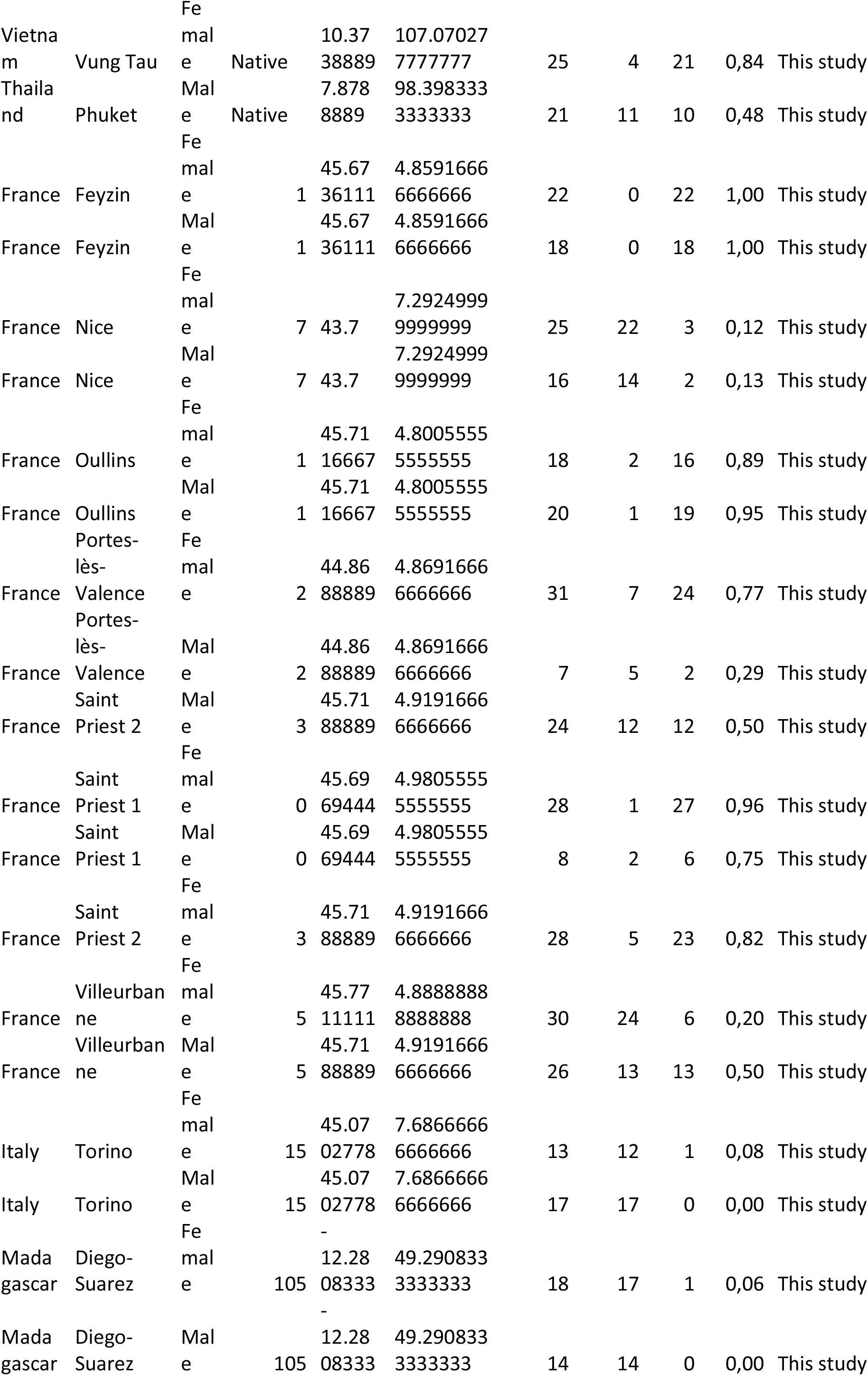
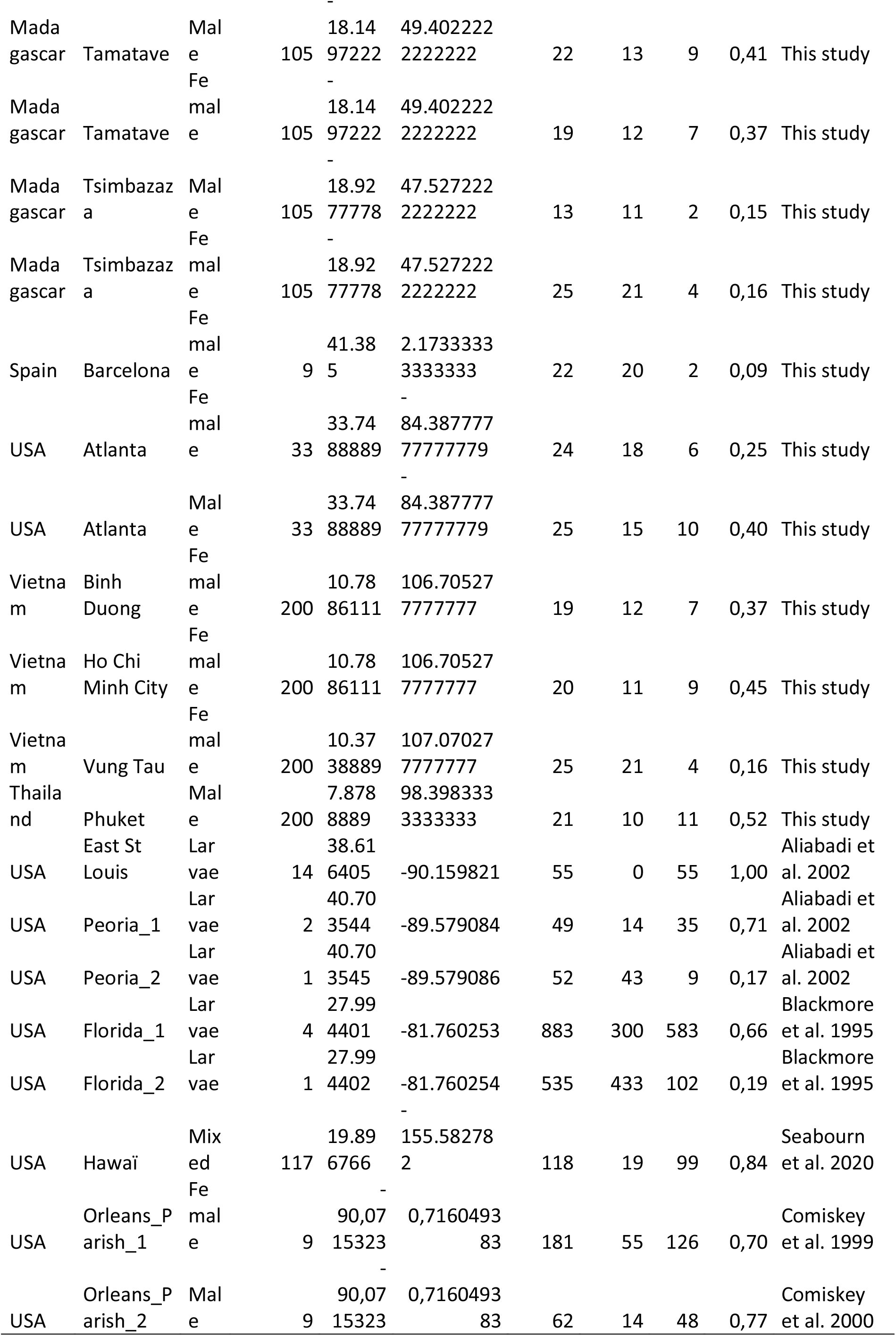
Number and details on mosquito prevalence from this study and the previously published studies.

## References

1. Atwoli L, Baqui AH, Benfield T, Bosurgi R, Godlee F, Hancocks S, et al. Call for Emergency Action to Limit Global Temperature Increases, Restore Biodiversity, and Protect Health: Wealthy Nations Must do Much More, Much Faster. Glob Heart. 2021;16(1):60.

2. Bellard C, Genovesi P, Jeschke JM. Global patterns in threats to vertebrates by biological invasions. Proceedings of the Royal Society B: Biological Sciences. 27 janv 2016;283(1823):20152454.

3. Bradshaw CJA, Leroy B, Bellard C, Roiz D, Albert C, Fournier A, et al. Massive yet grossly underestimated global costs of invasive insects. Nature Communications. 4 oct 2016;7:12986.

4. Didham RK, Tylianakis JM, Hutchison MA, Ewers RM, Gemmell NJ. Are invasive species the drivers of ecological change? Trends in Ecology & Evolution. 1 sept 2005;20(9):470-4.

5. Doherty TS, Glen AS, Nimmo DG, Ritchie EG, Dickman CR. Invasive predators and global biodiversity loss. Proc Natl Acad Sci U S A. 4 oct 2016;113(40):11261-5.

6. Mainka SA, Howard GW. Climate change and invasive species: double jeopardy. Integrative Zoology. 2010;5(2):102-11.

7. Hulme PE. Unwelcome exchange: International trade as a direct and indirect driver of biological invasions worldwide. One Earth. 21 mai 2021;4(5):666-79.

8. Andersen MC, Adams H, Hope B, Powell M. Risk Assessment for Invasive Species. Risk Analysis. 2004;24(4):787-93.

9. Diagne C, Leroy B, Vaissière AC, Gozlan RE, Roiz D, Jarić I, et al. High and rising economic costs of biological invasions worldwide. Nature. avr 2021;592(7855):571-6.

10. Hulme PE. Invasive species challenge the global response to emerging diseases. Trends in Parasitology. 1 juin 2014;30(6):267-70.

11. Shackleton RT, Shackleton CM, Kull CA. The role of invasive alien species in shaping local livelihoods and human well-being: A review. J Environ Manage. 1 janv 2019;229:145-57.

12. Chalkowski K, Lepczyk CA, Zohdy S. Parasite Ecology of Invasive Species: Conceptual Framework and New Hypotheses. Trends Parasitol. août 2018;34(8):655-63.

13. Roberts M, Dobson A, Restif O, Wells K. Challenges in modelling the dynamics of infectious diseases at the wildlife–human interface. Epidemics. 1 déc 2021;37:100523.

14. Dunn AM. Chapter 7 Parasites and Biological Invasions. In: Advances in Parasitology [Internet]. Academic Press; 2009 [cité 26 juin 2023]. p. 161-84. (Natural History of Host-Parasite Interactions; vol. 68).

15. Torchin ME, Lafferty KD, Kuris AM. Parasites and marine invasions. Parasitology. oct 2002;124(7):137-51.

16. Colautti RI, Ricciardi A, Grigorovich IA, MacIsaac HJ. Is invasion success explained by the enemy release hypothesis? Ecology Letters. 2004;7(8):721-33.

17. Drake JM. The paradox of the parasites: implications for biological invasion. Proceedings of the Royal Society of London Series B: Biological Sciences. 7 nov 2003;270(suppl_2):S133-5.

18. Blakeslee AMH, Fowler AE, Keogh CL. Chapter Two - Marine Invasions and Parasite Escape: Updates and New Perspectives. In: Lesser M, éditeur. Advances in Marine Biology [Internet]. Academic Press; 2013 [cité 26 juin 2023]. p. 87-169.

19. Llopis-Belenguer C, Blasco-Costa I, Balbuena JA, Sarabeev V, Stouffer DB. Native and invasive hosts play different roles in host–parasite networks. Ecography. 2020;43(4):559-68.

20. Llaberia-Robledillo M, Balbuena JA, Sarabeev V, Llopis-Belenguer C. Changes in native and introduced host–parasite networks. Biol Invasions. 1 févr 2022;24(2):543-55.

21. Nørgaard LS, Zilio G, Saade C, Gougat-Barbera C, Hall MD, Fronhofer EA, et al. An evolutionary trade-off between parasite virulence and dispersal at experimental invasion fronts. Ecol Lett. avr 2021;24(4):739-50.

22. Wilson K, Cotter S (2008) Density-dependent prophylaxis in insects. In: Ananthakrishnan TN, Whitman TW (eds) Phenotypic plasticity of insects: mechanisms and consequences, vol 1. Science Publishers, Enfeld, UK. 381–420.

23. Bradley CA, Altizer S. Parasites hinder monarch butterfly flight: implications for disease spread in migratory hosts. Ecology Letters. 2005;8(3):290-300.

24. Bonizzoni M, Gasperi G, Chen X, James AA. The invasive mosquito species Aedes albopictus: current knowledge and future perspectives. Trends Parasitol. sept 2013;29(9):460-8.

25. Lequime S, Dehecq JS, Matheus S, de Laval F, Almeras L, Briolant S, et al. Modeling intra-mosquito dynamics of Zika virus and its dose-dependence confirms the low epidemic potential of Aedes albopictus. PLoS Pathog. 31 déc 2020;16(12):e1009068.

26. Paupy C, Delatte H, Bagny L, Corbel V, Fontenille D. Aedes albopictus, an arbovirus vector: From the darkness to the light. Microbes and Infection. 1 déc 2009;11(14):1177-85.

27. Goubert C, Minard G, Vieira C, Boulesteix M. Population genetics of the Asian tiger mosquito Aedes albopictus, an invasive vector of human diseases. Heredity (Edinb). sept 2016;117(3):125-34.

28. Bennett K, Gómez Martínez C, Almanza A, Rovira JR, McMillan WO, Enriquez V, et al. High infestation of invasive Aedes mosquitoes in used tires along the local transport network of Panama. Parasites Vectors. déc 2019;12(1):1-10.

29. Eritja R, Palmer JRB, Roiz D, Sanpera-Calbet I, Bartumeus F. Direct Evidence of Adult Aedes albopictus Dispersal by Car. Sci Rep. 24 oct 2017;7(1):14399.

30. Madon MB, Mulla MS, Shaw MW, Kluh S, Hazelrigg JE. Introduction of Aedes albopictus (Skuse) in southern California and potential for its establishment. J Vector Ecol. juin 2002;27(1):149-54.

31. Medley KA, Jenkins DG, Hoffman EA. Human-aided and natural dispersal drive gene flow across the range of an invasive mosquito. Mol Ecol. 1 janv 2015;24(2):284-95.

32. Miller MJ, Loaiza JR. Geographic Expansion of the Invasive Mosquito Aedes albopictus across Panama—Implications for Control of Dengue and Chikungunya Viruses. PLOS Neglected Tropical Diseases. 8 janv 2015;9(1):e0003383.

33. Schmidt TL, Swan T, Chung J, Karl S, Demok S, Yang Q, et al. Spatial population genomics of a recent mosquito invasion. Molecular Ecology. 2021;30(5):1174-89.

34. Cornel AJ, Hunt RH. Aedes albopictus in Africa? First records of live specimens in imported tires in Cape Town. Journal of the American Mosquito Control Association. 1991;7(1):107-8.

35. Dalla Pozza G, Majori G. First record of Aedes albopictus establishment in Italy. J Am Mosq Control Assoc. sept 1992;8(3):318-20.

36. Laird M, Calder L, Thornton RC, Syme R, Holder PW, Mogi M. Japanese Aedes albopictus among four mosquito species reaching New Zealand in used tires. J Am Mosq Control Assoc. mars 1994;10(1):14-23.

37. Moore CG, Mitchell CJ. Aedes albopictus in the United States: ten-year presence and public health implications. Emerging Infect Dis. sept 1997;3(3):329-34.

38. Pluskota B, Storch V, Braunbeck T, Beck M, Becker N. First record of Stegomyia albopicta (Skuse) (Diptera: Culicidae) in Germany. European Mosquito Bulletin. 2008;26:1-5.

39. Schaffner F, Van Bortel W, Coosemans M. First record of Aedes (Stegomyia) albopictus in Belgium. J Am Mosq Control Assoc. juin 2004;20(2):201-3.

40. Juliano SA, O’Meara GF, Morrill JR, Cutwa MM. Desiccation and thermal tolerance of eggs and the coexistence of competing mosquitoes. Oecologia. 1 févr 2002;130(3):458-69.

41. Toma L, Severini F, Di Luca M, Bella A, Romi R. Seasonal patterns of oviposition and egg hatching rate of Aedes albopictus in Rome. J Am Mosq Control Assoc. mars 2003;19(1):19-22.

42. Sherpa S, Renaud J, Guéguen M, Besnard G, Mouyon L, Rey D, et al. Landscape does matter: Disentangling founder effects from natural and human-aided post-introduction dispersal during an ongoing biological invasion. Journal of Animal Ecology. 2020;89(9):2027-42.

43. Kambhampati S, Black WC 4th, Rai KS. Geographic origin of the US and Brazilian Aedes albopictus inferred from allozyme analysis. Heredity (Edinb). août 1991;67 (Pt 1):85-93.

44. Minard G, Tran FH, Van VT, Goubert C, Bellet C, Lambert G, et al. French invasive Asian tiger mosquito populations harbor reduced bacterial microbiota and genetic diversity compared to Vietnamese autochthonous relatives. Front Microbiol [Internet]. 22 sept 2015.

45. Gojković N, Ludoški J, Krtinić B, Milankov V. The First Molecular and Phenotypic Characterization of the Invasive Population of Aedes albopictus (Diptera: Culicidae) from the Central Balkans. J Med Entomol. 3 sept 2019;56(5):1433-40.

46. Schmidt TL, Chung J, Honnen AC, Weeks AR, Hoffmann AA. Population genomics of two invasive mosquitoes (Aedes aegypti and Aedes albopictus) from the Indo-Pacific. PLoS Negl Trop Dis. juill 2020;14(7):e0008463.

47. Vavassori L, Saddler A, Müller P. Active dispersal of Aedes albopictus: a mark-release-recapture study using self-marking units. Parasites Vectors. déc 2019;12(1):1-14.

48. Thongsripong P, Chandler JA, Green AB, Kittayapong P, Wilcox BA, Kapan DD, et al. Mosquito vector-associated microbiota: Metabarcoding bacteria and eukaryotic symbionts across habitat types in Thailand endemic for dengue and other arthropod-borne diseases. Ecol Evol. 2018;8(2):1352-68.

49. Lantova L, Volf P. Mosquito and sand fly gregarines of the genus Ascogregarina and Psychodiella (Apicomplexa: Eugregarinorida, Aseptatorina) - Overview of their taxonomy, life cycle, host specificity and pathogenicity. Infect Genet Evol. 4 mai 2014.

50. Erthal JA, Soghigian JS, Livdahl T. Life Cycle Completion of Parasite Ascogregarina taiwanensis (Apicomplexa: Lecudinidae) in Non-Native Host Ochlerotatus japonicus (Diptera: Culicidae). Journal of Medical Entomology. 1 sept 2012;49(5):1109-17.

51. Garcia JJ, Fukuda T, Becnel JJ. Seasonality, prevalence and pathogenicity of the gregarine Ascogregarina taiwanensis (Apicomplexa: Lecudinidae) in mosquitoes from Florida. J Am Mosq Control Assoc. sept 1994;10(3):413-8.

52. Levine ND. Progress in taxonomy of the Apicomplexan protozoa. J Protozool. nov 1988;35(4):518-20.

53. Munstermann LE, Wesson DM. First record of Ascogregarina taiwanensis (Apicomplexa: Lecudinidae) in North American Aedes albopictus. J Am Mosq Control Assoc. juin 1990;6(2):235-43.

54. Reeves WK, McCullough SD. Laboratory susceptibility of Wyeomyia smithii (Diptera: Culicidae) to Ascogregarina taiwanensis (Apicomplexa: Lecudinidae). J Eukaryot Microbiol. oct 2002;49(5):391-2.

55. Tseng M. Ascogregarine parasites as possible biocontrol agents of mosquitoes. J Am Mosq Control Assoc. 2007;23(2 Suppl):30-4.

56. Boisard J, Florent I. Why the –omic future of Apicomplexa should include gregarines. Biology of the Cell. 2020;112(6):173-85.

57. Aliabadi BW, Juliano SA. Escape from gregarine parasites affects the competitive interactions of an invasive mosquito. Biol Invasions. 1 sept 2002;4(3):283-97.

58. Comiskey NM, Lowrie RC, Wesson DM. Effect of nutrient levels and Ascogregarina taiwanensis (Apicomplexa: Lecudinidae) infections on the vector competence of Aedes albopictus (Diptera: Culicidae) for Dirofilaria immitis (Filarioidea: Onchocercidae). J Med Entomol. janv 1999;36(1):55-61.

59. Soghigian J, Livdahl T. Differential response to mosquito host sex and parasite dosage suggest mixed dispersal strategies in the parasite Ascogregarina taiwanensis. PLOS ONE. 13 sept 2017;12(9):e0184573.

60. Blackmore MS, Scoles GA, Craig GB. Parasitism of Aedes aegypti and Ae. albopictus (Diptera: Culicidae) by Ascogregarina spp. (Apicomplexa: Lecudinidae) in Florida. J Med Entomol. nov 1995;32(6):847-52.

61. Rueda LM. Pictorial keys for the identification of mosquitoes (Diptera: Culicidae) associated with dengue virus transmission. 2004;(589):1-60.

62. Knudsen AB, Romi R, Majori G. Occurrence and spread in Italy of Aedes albopictus, with implications for its introduction into other parts of Europe. J Am Mosq Control Assoc. juin 1996;12(Pt 1):177-83.

63. Collantes F, Delacour S, Alarcón-Elbal PM, Ruiz-Arrondo I, Delgado JA, Torrell-Sorio A, et al. Review of ten-years presence of Aedes albopictus in Spain 2004–2014: known distribution and public health concerns. Parasites Vectors. déc 2015;8(1):1-11.

64. Womack ML, Thuma TS, Evans BR. Distribution of Aedes albopictus in Georgia, USA. J Am Mosq Control Assoc. juin 1995;11(2 Pt 1):237.

65. Edwards FW. Notes on the Mosquitos of Madagascar, Mauritius and Réunion. Bulletin of Entomological Research. août 1920;11(2):133-8.

66. Comiskey NM, Lowrie RC Jr, Wesson DM. Role of Habitat Components on the Dynamics of Aedes albopictus (Diptera: Culicidae) from New Orleans. Journal of Medical Entomology. 1 mai 1999;36(3):313-20.

67. Seabourn P, Spafford H, Yoneishi N, Medeiros M. The Aedes albopictus (Diptera: Culicidae) microbiome varies spatially and with Ascogregarine infection. PLoS Negl Trop Dis. août 2020;14(8):e0008615.

68. Schneider CA, Rasband WS, Eliceiri KW. NIH Image to ImageJ: 25 years of image analysis. Nat Methods. juill 2012;9(7):671-5.

69. Schloss PD. Secondary structure improves OTU assignments of 16S rRNA gene sequences. ISME J. mars 2013;7(3):457-60.

70. Clements AN. The Biology of Mosquitoes: Sensory reception and behaviour. Chapman & Hall; 1999. 764 p.

71. Flacio E, Engeler L, Tonolla M, Lüthy P, Patocchi N. Strategies of a thirteen year surveillance programme on Aedes albopictus (Stegomyia albopicta) in southern Switzerland. Parasites Vectors. déc 2015;8(1):1-18.

72. Ziegler R, Blanckenhorn WU, Mathis A, Verhulst NO. Video analysis of the locomotory behaviour of Aedes aegypti and Ae. japonicus mosquitoes under different temperature regimes in a laboratory setting. Journal of Thermal Biology. 1 avr 2022;105:103205.

73. R Core Team. R: A Language and Environment for Statistical Computing. R Foundation for Statistical Computing. 2016.

74. Vega-Rúa A, Marconcini M, Madec Y, Manni M, Carraretto D, Gomulski LM, et al. Vector competence of Aedes albopictus populations for chikungunya virus is shaped by their demographic history. Commun Biol. 24 juin 2020;3(1):1-13.

75. Sherpa S, Blum MGB, Capblancq T, Cumer T, Rioux D, Després L. Unraveling the invasion history of the Asian tiger mosquito in Europe. Mol Ecol. 8 mars 2019;

76. Schatz AM, Park AW. Patterns of host–parasite coinvasion promote enemy release and specialist parasite spillover. Journal of Animal Ecology. 2023;92(5):1029-41.

77. Roche B, Léger L, L’Ambert G, Lacour G, Foussadier R, Besnard G, et al. The Spread of Aedes albopictus in Metropolitan France: Contribution of Environmental Drivers and Human Activities and Predictions for a Near Future. PLOS ONE. 11 mai 2015;10(5):e0125600.

78. Juliano SA, Lounibos LP. Ecology of invasive mosquitoes: effects on resident species and on human health. Ecol Lett. mai 2005;8(5):558-74.

79. Miles HB. The Mode of Transmission of the Acephaline Gregarine Parasites of Earthworms. The Journal of Protozoology. 1962;9(3):303-6.

80. King BJ, Monis PT. Critical processes affecting Cryptosporidium oocyst survival in the environment. Parasitology. mars 2007;134(3):309-23.

81. Jenkins MC, Parker C, O’Brien C, Miska K, Fetterer R. Differing Susceptibilities of Eimeria acervulina, Eimeria maxima, and Eimeria tenella Oocysts to Desiccation. Journal of Parasitology. 1 oct 2013;99(5):899-902.

82. Janouškovec J, Paskerova GG, Miroliubova TS, Mikhailov KV, Birley T, Aleoshin VV, et al. Apicomplexan-like parasites are polyphyletic and widely but selectively dependent on cryptic plastid organelles. McCutcheon J, Weigel D, Howe C, McFadden G, éditeurs. eLife. 16 août 2019;8:e49662.

83. Arranz-Solís D, Warschkau D, Fabian BT, Seeber F, Saeij JPJ. Late Embryogenesis Abundant Proteins Contribute to the Resistance of Toxoplasma gondii Oocysts against Environmental Stresses. mBio. 21 févr 2023;14(2):e02868-22.

84. Bennett K, Gómez-Martínez C, Almanza A, Rovira J, McMillan W, Enriquez V, et al. High infestation of invasive Aedes mosquitoes in used tires along the local transport network of Panama. Parasites & Vectors. 27 mai 2019;12.

85. Schmidt TL, Rašić G, Zhang D, Zheng X, Xi Z, Hoffmann AA. Genome-wide SNPs reveal the drivers of gene flow in an urban population of the Asian Tiger Mosquito, Aedes albopictus. PLOS Neglected Tropical Diseases. 18 oct 2017;11(10):e0006009.

86. Dunn AM. Parasites and biological invasions. Adv Parasitol. 2009;68:161-84.

87. Buck JC, Hechinger RF, Wood AC, Stewart TE, Kuris AM, Lafferty KD. Host density increases parasite recruitment but decreases host risk in a snail–trematode system. Ecology. 2017;98(8):2029-38.

88. Watve MG, Sukumar R. Parasite abundance and diversity in mammals: correlates with host ecology. Proc Natl Acad Sci USA. 12 sept 1995;92(19):8945-9.

89. Ruiz-González MX, Moret Y, Brown MJF. Rapid induction of immune density-dependent prophylaxis in adult social insects. Biology Letters. 23 déc 2009;5(6):781-3.

90. Silva FWS, Serrão JE, Elliot SL. Density-dependent prophylaxis in primary anti-parasite barriers in the velvetbean caterpillar. Ecological Entomology. 2016;41(4):451-8.

91. Davis AK, Roode JC de. Effects of the parasite, Ophryocystis elektroscirrha, on wing characteristics important for migration in the monarch butterfly. Animal Migration. 1 déc 2018;5(1):84-93.

92. Marden JH, Cobb JR. Territorial and mating success of dragonflies that vary in muscle power output and presence of gregarine gut parasites. Animal Behaviour. oct 2004;68(4):857-65.

93. Schilder RJ, Marden JH. Parasites, proteomics and performance: effects of gregarine gut parasites on dragonfly flight muscle composition and function. Journal of Experimental Biology. 15 déc 2007;210(24):4298-306.

94. Girard M, Martin E, Vallon L, Raquin V, Bellet C, Rozier Y, et al. Microorganisms Associated with Mosquito Oviposition Sites: Implications for Habitat Selection and Insect Life Histories. Microorganisms. août 2021;9(8):1589.

95. Wooding M, Naudé Y, Rohwer E, Bouwer M. Controlling mosquitoes with semiochemicals: a review. Parasites & Vectors. 17 févr 2020;13(1):80.

96. Kaufmann C, Collins LF, Brown MR. Influence of Age and Nutritional Status on Flight Performance of the Asian Tiger Mosquito Aedes albopictus (Diptera: Culicidae). Insects. 26 juill 2013;4(3):404-12.

97. Stump E, Childs LM, Walker M. Parasitism of Aedes albopictus by Ascogregarina taiwanensis lowers its competitive ability against Aedes triseriatus. Parasites & Vectors. 25 janv 2021;14(1):79.

98. Comiskey NM, Lowrie RC, Wesson DM. Role of habitat components on the dynamics of Aedes albopictus (Diptera: Culicidae) from New Orleans. J Med Entomol. mai 1999;36(3):313-20.

99. Westby KM, Sweetman BM, Van Horn TR, Biro EG, Medley KA. Invasive species reduces parasite prevalence and neutralizes negative environmental effects on parasitism in a native mosquito. Journal of Animal Ecology. 2019;88(8):1215-25.

100. McIntire KM, Chappell KM, Juliano SA. How do noncompetent hosts cause dilution of parasitism? Testing hypotheses for native and invasive mosquitoes. Ecology. oct 2021;102(10):e03452.

101. Dos Passos RA, Tadei WP. Parasitism of Ascogregarina taiwanensis and Ascogregarina culicis (Apicomplexa: Lecudinidae) in larvae of Aedes albopictus and Aedes aegypti (Diptera: Culicidae) from Manaus, Amazon region, Brazil. J Invertebr Pathol. mars 2008;97(3):230-6.

102. Bagny Beilhe L, Delatte H, Juliano SA, Fontenille D, Quilici S. Ecological interactions in Aedes species on Reunion Island. Medical and Veterinary Entomology. 2013;27(4):387-97.

103. Costanzo KS, Mormann K, Juliano SA. Asymmetrical Competition and Patterns of Abundance of Aedes albopictus and Culex pipiens (Diptera: Culicidae). Journal of Medical Entomology. 1 juill 2005;42(4):559-70.

104. de Oliveira S, Villela DAM, Dias FBS, Moreira LA, Maciel de Freitas R. How does competition among wild type mosquitoes influence the performance of Aedes aegypti and dissemination of Wolbachia pipientis? PLoS Negl Trop Dis. 9 oct 2017;11(10):e0005947.

105. Juliano SA. Species Introduction and Replacement Among Mosquitoes: Interspecific Resource Competition or Apparent Competition? Ecology. 1998;79(1):255-68.

106. Kesavaraju B, Leisnham PT, Keane S, Delisi N, Pozatti R. Interspecific Competition between Aedes albopictus and A. sierrensis: Potential for Competitive Displacement in the Western United States. PLoS ONE. 28 févr 2014;9(2):e89698.

107. O’Donnell DL, Armbruster P. Comparison of Larval Foraging Behavior of Aedes albopictus and Aedes japonicus (Diptera: Culicidae). Journal of Medical Entomology. 1 nov 2007;44(6):984-9.

